# Distinct Programs Drive Organ Development and Regeneration

**DOI:** 10.1101/2025.11.20.689429

**Authors:** Rongze Ma, ZhiXuan Deng, Yiting Li, Jing Chen, Jiayi Zhang, Deyi Feng, Chongshen Xu, Jiecan Zhou, Xiaodong Zhang, Kun Xia, Pengfei Lu

## Abstract

A prevailing paradigm in stem cell and developmental biology posits that regeneration is a recapitulation of embryonic programs, yet the molecular logic underlying these processes remains unclear. Here, using the mammary gland as a model, we identified a novel intermediate cell state essential for basal-to-luminal differentiation during regeneration. The transcription factor *Pou2f3* is a master regulator of this state, critical for regeneration but dispensable for organ physiology. Mechanistically, we found that POU2F3 functions as a transcriptional repressor that drives lineage conversion by restricting chromatin accessibility of basal genes rather than by directly activating luminal programs. Furthermore, we uncovered a POU2F3-TNF feedback loop that maintains epithelial homeostasis, where TNF signaling suppresses *Pou2f3* to inhibit differentiation, and loss of luminal cells relieves this repression to trigger regeneration. Strikingly, cell fate is governed by signaling pathways essential for cell-cell communication during development, but by *Pou2f3*-dependent chromatin remodeling for rapid response and injury repair. We validate the regeneration-specific requirement for *Pou2f3* in mammary gland and further demonstrate analogous regeneration defects in prostate and pancreas. These findings challenge the long-held assumption that regeneration recapitulates development and establish a framework for understanding context-specific mechanisms of tissue plasticity, with implications for regenerative medicine and epithelial biology.

## INTRODUCTION

A central question in biology is understanding how a single cell gives rise to a vast cellular diversity through differentiation and builds a three-dimensional architecture through morphogenesis to sustain the physiology of a functional organ. For more than sixty years, a foundational dogma in developmental and regenerative biology has been that tissue regeneration is a process that recapitulates the embryonic developmental programs that initially built the organ ^1^. This is indeed supported by decades of subsequent studies showing that the processes of embryonic development, tissue homeostasis, and regeneration all rely on similar decisions that guide stem cells and progenitors toward their final, differentiated fates^2,3^. However, the precise molecular programs driving cell fate decisions in these two distinct biological contexts—organ development and adult regeneration—remain poorly understood in vertebrate organs ^4^.

As a well-characterized vertebrate organ, the mouse mammary gland is an ideal model for understanding cell fate decisions in developmental and regenerative biology. As an ectodermal derivative, mammary gland development starts relatively late compared to most other organs. Embryonic mammary stem cells initially expressing keratin 14 (K14) are generally considered to be specified around embryonic day (E) 13.5 to E15.5. They eventually mature into K14^+^ adult basal cells and K8^+^ luminal cells, which, depending on their expression of either estrogen receptor (ER) or ELF5, can be divided into hormone-responsive luminal cells (HRLCs) and secretory luminal cells (SLCs), respectively (Fig. 1A) ^5^. Various signaling pathways, including WNT, Notch, and Integrin signaling, govern these differentiation events and concurrent morphogenetic processes, e.g. epithelial branching ^6,7^. Although the first branch forms in embryogenesis, the initial epithelial network at birth is very primitive and remains relatively dormant until puberty when surging female hormones initiate vigorous branching until the stromal fat-pad is filled by the epithelium ^8^. In adulthood, each pregnancy induces a cycle of alveolar expansion, lactation, and involution, highlighting dynamic fate plasticity ^9^.

**Figure 1:**
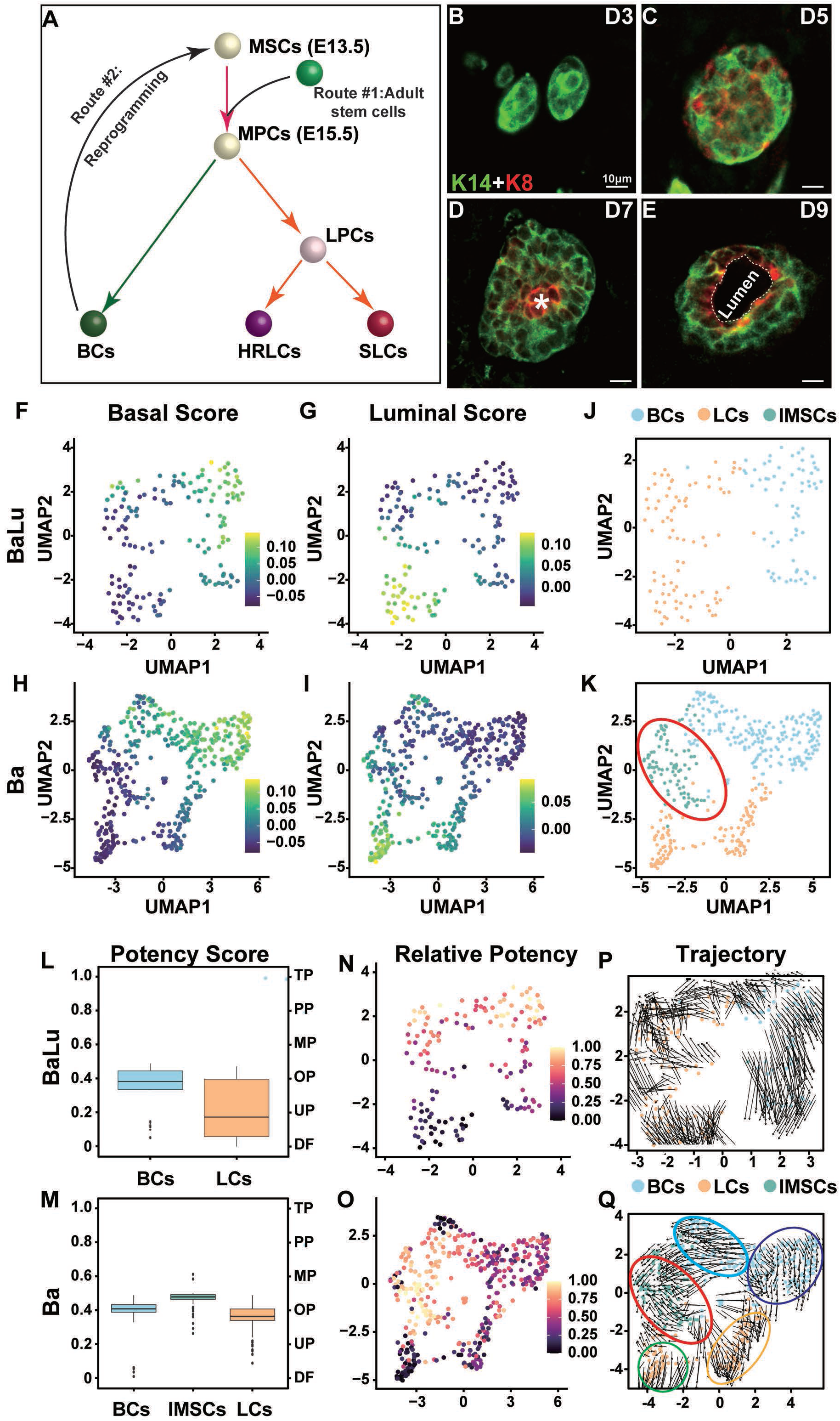
Identification of a Novel Intermediate Cell State During Mammary Gland Regeneration. (**A**) Diagram of the current model of mammary gland stem cell differentiation during development and regeneration. Abbreviations: MSCs, mammary stem cells; MPCs, mammary progenitor cells; BCs, basal cells; HRLCs, hormone-responsive luminal cells; SLCs, secretory luminal cells. (**B-E**) Immunofluorescence staining of the basal cell marker K14 (green) and the luminal cell marker K8 (red) in mammary glands at days 3, 5, 7, and 9 post-transplantation of purified basal cells into cleared fat pads of immunodeficient mice. Scale bars: 10 µm. (**F-K**) Single-cell RNA sequencing (scRNA-seq) analysis of transplants derived from a mixture of basal and luminal cells (BaLu) and purified basal cells alone (Ba) at day 5 post-transplantation. Dimensionality reduction and marker-based classification revealed distinct basal and luminal populations in both conditions. UMAP plot showing cell clusters (**F-I**) and expression of basal, luminal, and intermediate cell markers (**J-K**). An intermediate state cells (IMSCs), characterized by low expression of both basal and luminal markers, was identified in the Ba samples but was absent in the BaLu samples. Abbreviations: BCs, basal cells; LCs, luminal cells; IMSCs, intermediate state cells. (**L-O**) Cytotrace analysis showing stemness scores of intermediate, basal, and luminal cells. Intermediate cells exhibited higher stemness scores than basal and luminal cells, indicating a more plastic state. (**L, M**) Comparison of stemness scores between cell populations. (**N, O**) UMAP plots showing the distribution of stemness scores. (**P, Q**) RNA velocity analysis revealing the differentiation trajectory. BaLu samples (**P**) exhibited minimal differentiation dynamics. By contrast, basal cell subgroup 1(BCS1; blue ellipse) in the Ba samples (**Q**) showed a clear trajectory from basal cells transitioning through the intermediate state (red ellipse) toward HRLC (green circle) and SLC (orange ellipse) differentiation, while the rest basal cell subgroup 2 (BCS2, dark blue ellipse) showed little differentiation dynamics. Annotations on the plot indicate cell potency: TP, totipotency; PP, pluripotency; MP, multipotency; OP, oligopotency; UP, unipotency; DF, differentiated.

The mammary gland also has a long history of being used as a model to study stem cell biology since it was shown that a piece of the epithelium could be repeatedly transplanted into a “cleared” fat-pad free of its endogenous epithelium and regenerate a functional gland ^10^. However, subsequent studies showed that luminal and basal cells remain committed during mammary gland regeneration from a piece of the epithelium ^11,12^. This is similar to regeneration of the axolotl limbs ^13,14^, fish fins ^15^, and mouse digits ^16^ where lineage-committed cells proliferate and replenish the missing tissue. By contrast, when the luminal population is under-represented^11,17^, for example in a transplantation model involving only basal cells, a small portion of basal cells, especially those expressing *Lgr5*, *Procr*, *Tspan*, etc., exhibit multiplicity and give rise to luminal cells ^18–20^. Further studies show that basal multiplicity is negatively regulated the levels of luminal-secreted TNF, although its mechanistic details remain elusive at present ^21,22^.

At present, how the *Lgr5*^+^ or *Procr*^+^ cells acquire their “stemness” or multiplicity—whether it is an intrinsic property of a rare quiescent population of adult stem cells or an extrinsic property acquired during the transplantation process—remains a topic of active debate^6,23^ (Fig. 1A). Despite their difference in opinion, both schools of thought agree that basal stem cells follow the same developmental program, e.g. through a transient intermediate, or “hybrid” state expressing both basal and luminal markers, to give rise to luminal cells ^21,22^ (Fig. 1A). Consistent with this belief, various pathways, including Notch and Integrin signaling, govern luminal differentiation in both development and regeneration based on basal cells^24^. Despite the above framework, the molecular underpinnings of cell fate decisions in mammary gland development and regeneration remain largely unclear. To address this question, we searched for transitional cell states during basal-to-luminal conversion and characterized candidate regulators using single-cell transcriptomics, epigenomics and functional genetics.

## RESULTS

### Identification of a Novel Intermediate Cell State During Mammary Gland Regeneration

To elucidate the differentiation mechanism of basal cells during mammary gland regeneration, we first asked whether there is a distinct intermediate stage during basal-to-luminal conversion in regeneration. Therefore, we used fluorescence-assisted cell sorting (FACS) to isolate basal cells (CD49f^high^CD24^med^) from dissociated mammary glands of the *R26R*^mTmG^ mouse line ^25^. Ten thousand purified basal cells were then transplanted into the cleared fat pads of immunodeficient nude mice and the transplants were harvested on days 3, 5, 7, and 9 after surgery. They were then frozen-sectioned, stained for the basal marker K14 and luminal marker K8, and examined by confocal microscopy.

We found that transplanted basal cells did not express the luminal marker K8 until day 5 (Fig. 1B, C). At this stage, however, all basal cells still expressed K14, and those that expressed K8 were double-positive for both (Fig. 1C). By day 7, the K8^+^ cells, which were distributed throughout the aggregate (Fig. 1D), had converged to the center and lost K14 expression. By day 9, a lumen lined by K8-positive cells began to emerge, with the K14^+^ basal cells positioned farthest from the lumen in the epithelial aggregate (Fig. 1E). Because the structure and architecture of the growth resembled a developing epithelial duct, we concluded that mature luminal cells had been generated from the transplanted basal cells. We considered day 5 a putative intermediate stage because many transplanted cells co-expressed K14 and K8, indicative of a transitional identity.

To test this, we performed scRNA-seq to determine the transcriptional changes of transplanted basal cells on day 5 (Ba sample). As a control, we also performed scRNA-seq on samples derived from the transplantation of mammary epithelial cell aggregates with basal and luminal cells mixed at a 1:1 ratio (BaLu sample). After clustering and dimensionality reduction, we scored the cells using previously published markers for basal and luminal cells^26^ (Fig. 1F-K, Supplementary Fig. 1A, B). Interestingly, similar to the BaLu group, we also observed distinct, seemingly mature basal and luminal populations in the Ba group. The presence of mature luminal cells based on mRNA but not on protein (Fig. 1E) expression is presumably a result of their different kinetics and half-lives during luminal maturation. Moreover, we observed a unique cell population in the Ba group that exhibited intermediate levels of both basal and luminal markers, and this population was absent in the BaLu samples (Fig. 1J-K, Supplementary Fig. 1A, B). We propose that this population represents an intermediate cell state during differentiation.

Using CytoTRACE ^27^, we scored the differentiation potential of cells of the transplanted samples. We found that the intermediate cells exhibited a higher stemness score compared to both the basal and luminal cells, whereas such a highly stem-like subpopulation was not observed in the BaLu samples (Fig. 1L-O). Furthermore, RNA velocity analysis ^28^ revealed a differentiation trajectory in the Ba samples where a subgroup of basal cells, which we called basal cell subgroup 1 (BCS1) progressed toward luminal cells via the intermediate state. By contrast, another subgroup of basal cells in the Ba samples showed minimal differentiation trajectory, which we referred to as basal cell subgroup 2 (BCS2). Similar to BCS2 cells, cells in the BaLu samples primarily progressed within their own subpopulations (Fig. 1P, Q). Moreover, RNA velocity analysis also indicated that the intermediate state cells generated *ER*^+^ HRLCs and *Elf5*^+^ SLCs in parallel (Fig. 1Q, Supplementary Fig. 1C). We also found that this intermediate cell population was highly proliferative, with nearly all cells in the G2/M and S phases (Supplementary Fig. 1D, E).

Finally, when we scored our data with the marker genes for the “hybrid-cell” (CD29^high^EpCAM^high^) previously proposed to mediate basal-to-luminal conversion^21,22^, we found that they could not reliably identify the intermediate cell population in our samples (Supplementary Fig. 1F, G), suggesting that the IMSCs are distinct from these previously identified hybrid cells. Interestingly, previously reported stem cell markers such as *Tspan8*, *Procr*, *Bcl11b*, and *Lgr5* were not significantly enriched in the BSC1 or the IMSC populations (Supplementary Fig. 1H-K), despite the observation that basal cells expressing them are more likely to regenerate the mammary gland than those non-expressing cells ^18–20^.

In summary, our findings establish the presence of a transient intermediate cell state during mammary gland regeneration, characterized by increased proliferative capacity and a distinct transcriptional signature.

### The POU Domain Transcription Factor *Pou2f3* is a Key Regulator of the Intermediate State

The observation of this transient double-positive state at day 5 strongly suggests it represents a crucial intermediate stage in the basal-to-luminal conversion process. Genes uniquely expressed during this intermediate phase are thus strong candidates for regulating this lineage transition. To interrogate the mechanisms governing the basal-to-luminal cell state transition, we compared gene expression in the intermediate state cells with that in basal and luminal cells (Fig. 2A, Supplementary Fig. 2A). Gene Ontology (GO) enrichment analysis of upregulated pathways in the intermediate cells highlighted a profound shift towards proliferative and genomic activities, including chromosome segregation, DNA replication, and nuclear division (Fig. 2B). This enrichment in cell cycle and DNA replication pathways is consistent with a highly proliferative state necessary for tissue regeneration. Conversely, pathways downregulated in the intermediate cells reflect a reduction in extracellular matrix interactions, including ECM organization and cell-substrate adhesion (Supplementary Fig. 2B). The decrease in ECM and adhesion-related pathways suggests a transient detachment or remodeling of the cellular environment, facilitating the re-patterning required for luminal differentiation.

**Figure 2:**
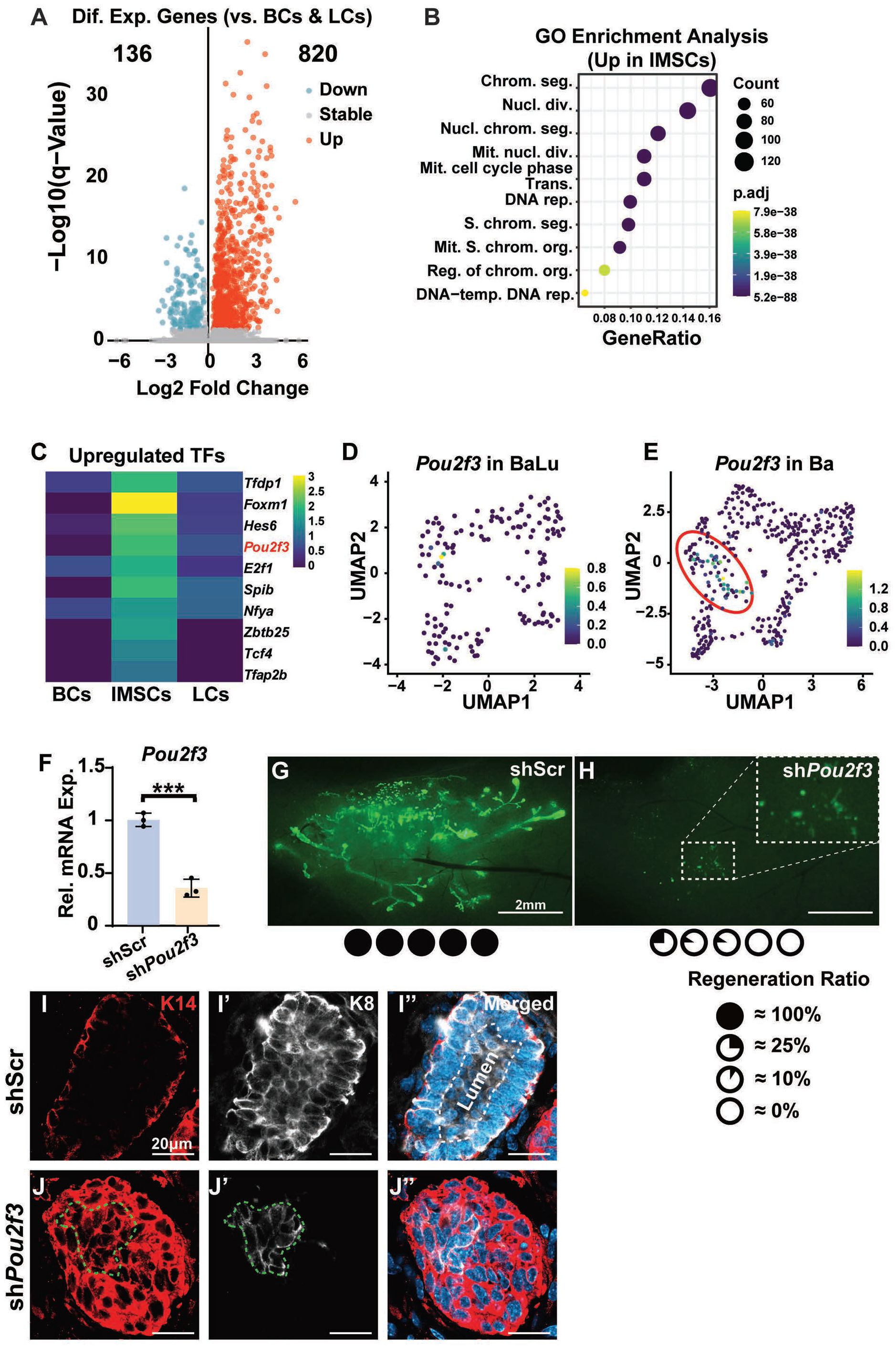
The POU Domain Transcription Factor *Pou2f3* is a Key Regulator of the Intermediate State. (**A**) Differentially expressed genes by intermediate state cells (IMSCs) when compared with basal cells (BCs) and luminal cells (LCs). (**B**) Gene ontology (GO) enrichment analysis of the upregulated genes in IMSCs, emphasizing pathways linked to rapid proliferation, chromosome remodeling, and differentiation. (**C**) Heatmap showing the SCENIC transcription factor activity analysis scores for the top upregulated transcription factors in IMSCs. *Pou2f3* was identified as a candidate regulator of luminal differentiation during regeneration. (**D, E**) *Pou2f3* expression in single cells of the UMAP plots from the BaLu (D) and Ba (E) samples. Note that *Pou2f3* expressing cells overlap well with IMSCs in the Ba samples. (**F-J**) Functional validation of *Pou2f3* in mammary gland regeneration. (F) qPCR analysis showing the knockdown efficiency of *Pou2f3* using shRNA. N=3. (**G**, **H**) Green-fluorescent images showing the epithelial outgrowths of the control shScr-infected (**G**) and *Pou2f3*-knockdown sh*Pou2f3*-infected (**H**) whole-mount glands. Inset shows that sh*Pou2f3* knockdown cells formed aggregates but failed to undergo epithelial branching as in controls. N=5. (**I**, **J**) Immunofluorescence staining showing reduced luminal cell differentiation in *Pou2f3*-deficient cells. Note that, contrary to shScr-samples, in which K14^+^ (red) basal and K8^+^ (white) luminal cells separated and formed a lumen, no mature luminal cells expressing only K8 (encircled by a green-dotted line) were found in the sh*Pou2f3* samples, and the mutant aggregate did not form a lumen (encircled by a white-dotted line). Data are mean ± SD. Statistical analysis was performed using unpaired Student’s t-test. *p < 0.05; **p < 0.01; ***p < 0.001; ****p < 0.0001; n.s., not significant.

To select a candidate regulator for functional testing, we decided to shorten the candidate list by focusing on the transcriptional factors potentially governing basal cell differentiation. Therefore, we performed SCENIC analysis on the Ba samples and identified a series of significantly activated transcription factors in the intermediate cells (Fig. 2C). Most of these candidate transcription factors were not specific to the intermediate cell state, except for *Pou2f3* and *Spib* that were barely expressed in the Balu samples (Supplementary Fig. 2C, D; Fig. 2D, E). Therefore, we decided to focus on the class II POU domain transcription factor *Pou2f3*, known to be a key regulator of the development of chemosensory tuft cells in mucosal epithelial tissues ^29,30^.

Next, we tested whether *Pou2f3* is required for the basal-to-luminal transition in vivo. Using shRNA, we reduced *Pou2f3* expression by ∼60%, as validated by qPCR (Fig. 2F), and transplanted these cells into cleared fat pads. In contrast to control shRNA-expressing glands (Fig. 2G), *Pou2f3* knockdown severely impaired mammary regeneration, with growth confined to the injection site and a failure to form branches (Fig. 2H). Immunofluorescence staining for basal and luminal markers showed that the control group formed mature luminal cells and ductal structures (Fig. 2I-I’’), while the sh*Pou2f3* group failed to generate mature luminal cells as most K8^+^ cells still retained K14 expression, and no lumen was observed (Fig. 2J-J’’).

Overall, these findings establish *Pou2f3* as a key regulator of basal-to-luminal differentiation during mammary gland regeneration and show that it is required for basal cells to advance beyond the intermediate state to take on a mature luminal fate.

### POU2F3 Promotes Basal-to-Luminal Conversion by Restricting Chromatin Accessibility to the Basal Transcriptional Program

Given the critical role of *Pou2f3* in regeneration and the historical use of transplantation as a classic method to assess stem cell function, we predicted that *Pou2f3* is also essential for mammary gland development. Indeed, *Pou2f3* is expressed in both basal and luminal cells, albeit at a relatively low level when compared to its expression in the IMSCs based in scRNA-seq data from development and regeneration (Supplementary Fig. 3A, B). To test its role in development, we generated a *Pou2f3* knockout mouse (Supplementary Fig. 3C, D) and confirmed its absence of mRNA expression in the mammary epithelium via qPCR (Supplementary Fig. 3E). Surprisingly, the *Pou2f3-*deficient female mutant mice were able to nurse their pups, which had a similar survival rate and weight gain as the control pups (Supplementary Fig. 3F, G), despite our prediction that the mammary gland would fail to develop in the absence of the gene. An examination of the whole-mount mammary gland during lactation, using Carmine staining, revealed that the mutant glands were fully developed as the control (Supplementary Fig. 3H, I), and H&E staining of the histological sections further confirmed that the alveoli were filled with milk (Supplementary Fig. 3H’, I’).

To examine whether the phenotype we saw in mammary gland regeneration was a result of off-target effects of the shRNA knockdown, we repeated the transplantation experiments using *Pou2f3*^-/-^ basal cells. We found that *Pou2f3*^-/-^ basal cells were completely unable to regenerate the mammary gland when transplanted into the fat pad (Supplementary Fig. 3J, K), thus confirming the shRNA data was a result of *Pou2f3* deficiency, rather than off-target effects. These results present an apparent paradox: *Pou2f3* is required for regeneration but appears dispensable for normal gland physiology.

As a starting point to explain such a paradox, we attempted to interrogate its molecular function. However, the failure of mammary gland regeneration and a lack of regenerative growths by basal cells lacking *Pou2f3* was a serious technical limitation to understanding its function. To circumvent this limitation, we decided to take advantage of a recent culture protocol whereby single mammary basal stem cells could regenerate an entire functional mammary gland in vitro ^31^. Single mammary basal cells were isolated and cultured for seven days before they were embedded as an organoid. Under this condition, a control organoid started to branch by day 4 and its branches began to elongate by day 10, until a minigland formed by day 14 (Fig. 3A-A’’’). Interestingly, although *Pou2f3* null organoids grew in size somewhat similarly during the organoid and branching organoid stages, their branches seemed much shorter and stubbier than normal and failed to elongate. They appeared to be stuck at the branching organoid stage and failed to form a minigland (Fig. 3B-B’’’).

**Figure 3:**
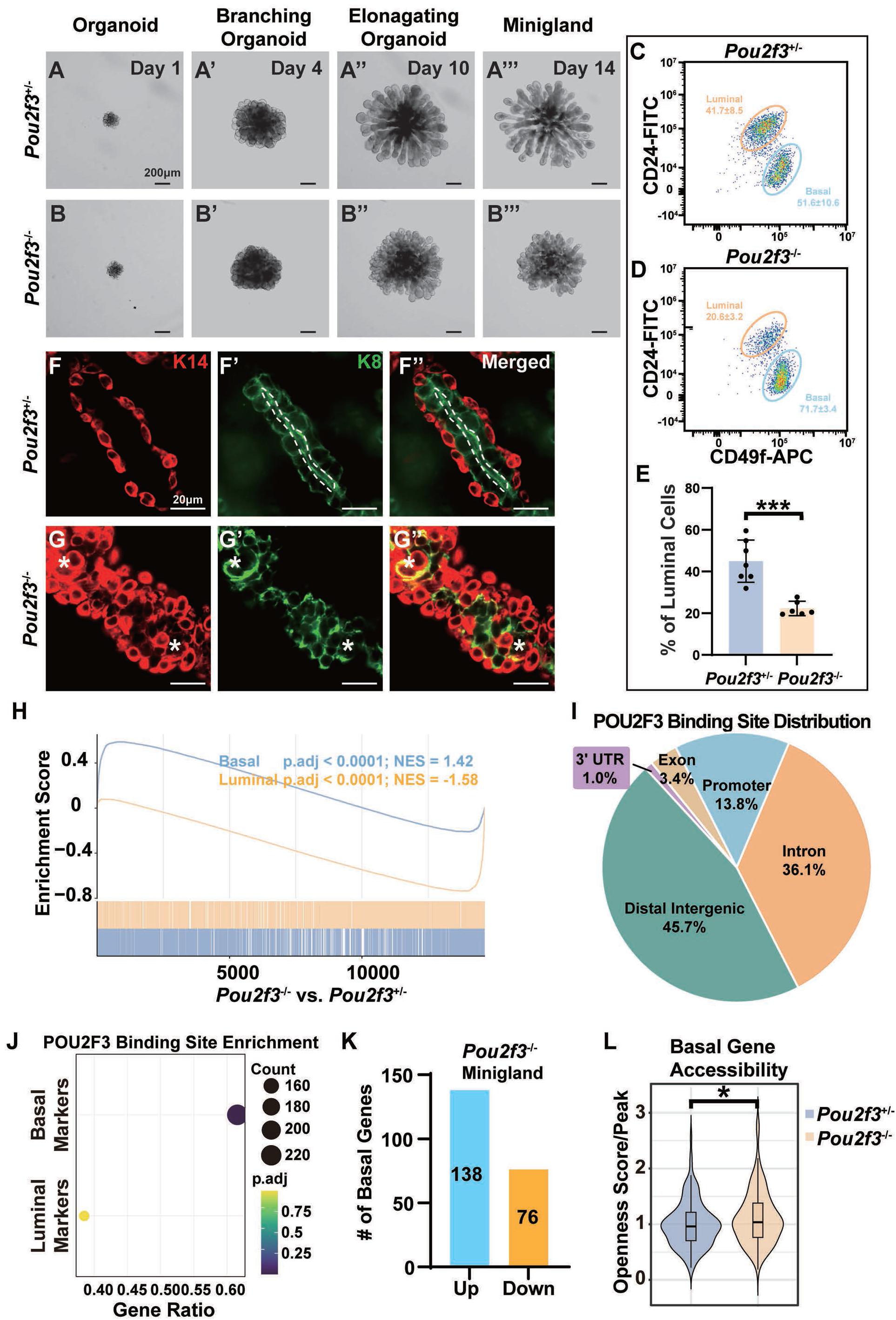
POU2F3 Promotes Basal-to-Luminal Conversion by Restricting Chromatin Accessibility to the Basal Transcriptional Program. (**A**-**E**) Characterization of mammary gland differentiation in vitro using a minigland culture assay. Control (*Pou2f3*^-/+^; **A**-**A’’’**) or *Pou2f3* null (**B**-**B’’’**) basal cells were embedded and cultured for 14 days. Representative images show the progression from the organoid (day 1), to branching organoid (day 4), elongating organoid (day 10), and minigland (day 14) stages. The effect of *Pou2f3* loss on luminal differentiation was quantified using FACS (**C**-**E**). N ≥ 6. (**F**-**G’’**) Immunofluorescent confocal microscopy of K14 (red) (F, G), K8 (green) (F’, G’), and merged (F’’, G’’) images of the miniglands derived from control (F-F’’) or *Pou2f3* null (G-G’’) basal cells. Asterisks indicate K14 and K8 double positive cells. N = 3. (**H**) GSEA analysis of basal and luminal programs in the miniglands derived from control and *Pou2f3* null basal cells. (**I**) Genomic distribution of POU2F3-binding sites in mammary epithelial cells based on CUT&Tag analysis. Note that binding sites were predominantly located in distal regulatory regions and intronic elements, suggesting a role in enhancer-mediated gene regulation. (**J**) Enrichment of POU2F3 binding regions of genes of basal and luminal marker genes based on CUT&Tag assays. (**K**) Bar chart showing the number of upregulated and downregulated basal genes bound by POU2F3. (**L**) Violin plot comparing the openness score of the chromatin of basal genes between control and null cells. Data are mean ± SD. Statistical analysis was performed using unpaired Student’s t-test. *p < 0.05; **p < 0.01; ***p < 0.001; ****p < 0.0001; n.s., not significant.

To assess the status of luminal differentiation, we first used flow cytometry based on the expression of CD49f and CD24 and analyzed the basal and luminal cell distribution of the dissociated single cells. Compared to the control, we found a 50.6% reduction in the percentage of luminal cells in the *Pou2f3* null epithelium compared to control (Fig. 3C-E). Next, we used immunofluorescence confocal microscopy and examined the expression of the basal marker K14 and luminal marker K8. As expected, we saw a double-layered mammary epithelial duct, with K14+ basal cells on the outside enveloping the K8+ luminal cells, with a mature lumen in the middle in the control organoids (Fig. 3F-F’’). This epithelial tissue architecture, however, was not observed in the *Pou2f3* null organoids. Specifically, despite the presence of many cells expressing exclusively K14 or K8, a lumen was never formed, and these cells also appeared misshapen. Luminal cells were round and piled up seemingly randomly, without the classic columnar shape indicative of epithelial architecture. We observed a few K14 and K8 double-positive cells, however, fewer than observed in vivo (Fig. 3G-G’’). Therefore, we concluded that the minigland culture is a relatively close approximation to, despite being a somewhat underestimate of, the in vivo regeneration assay.

To complement the loss-of-function (LOF) study based on the *Pou2f3* knockout strategy, we performed a gain-of-function (GOF) study where *Pou2f3* was forcefully expressed. Using a lentiviral overexpression construct, we found that *Pou2f3*-GOF led to a more than 6-fold increase of its mRNA expression (Supplementary Fig. 4A). Around half of the organoids failed to reach the branching organoid stage, with the rest showing branches that were longer and less dense than normal (Supplementary Fig. 4B-E). The branching abnormality is similar to that shown by luminal-derived organoids ^32^. Indeed, we saw a 72.5% increase in the percentage of luminal cells by flow cytometry in the *Pou2f3*-GOF organoids (Supplementary Fig. 4F-H). The GOF results are consistent with the LOF results, and, together, they show that *Pou2f3* is essential for the conversion of luminal cells from basal cells.

Using bulk RNA-sequencing, we found that basal signatures were upregulated, while luminal signatures were downregulated in *Pou2f3* null miniglands when compared to the control miniglands (Fig. 3H). In addition, an examination of the top 20 most differentially expressed genes in *Pou2f3* null miniglands showed downregulation of *Avil* (Supplementary Fig. 4I), which is consistent with it being a downstream target of *Pou2f3* in certain context ^33^. The reduced expression of genes like *Muc1* and *Kit* (Supplementary Fig. 4I), which are associated with epithelial differentiation and stem cell function respectively, aligns with the observed failure in luminal cell maturation.

We next sought to define the molecular mechanism by which POU2F3 regulates mammary gland cell fate. We hypothesized that POU2F3, as a transcription factor, directly controls gene expression. To test this, we performed the Cleavage Under Targets & Tagmentation (CUT&Tag) analysis to map the genome-wide binding sites of POU2F3 in basal cells from the minigland culture. We identified a large number of POU2F3-specific peaks, primarily located in distal intergenic regions (Fig. 3I), a distribution pattern consistent with previous findings in small cell lung cancer ^34^. Interestingly, the regulatory regions of a majority of the Notch genes, were not bound by POU2F3 (Supplementary Table 1), suggesting that Notch signaling activation is unlikely to be a direct consequence of POU2F3 transcriptional promotion. By contrast, basal cell-specific genes were significantly enriched among the genes bound by POU2F3, whereas luminal-specific genes were not (Fig. 3J), suggesting that POU2F3 specifically targets the transcriptional program of the basal cell lineage. Consistent with this notion, RNA-seq analysis showed that the expression of most bound basal genes was upregulated after *Pou2f3* knockout compared to control (Fig. 3K), indicating that POU2F3 primarily acts as a transcriptional repressor for most of these basal target genes.

Next, we performed ATAC-seq to assess chromatin accessibility in control and *Pou2f3* knockout miniglands. We found that the chromatin accessibility at the POU2F3-bound basal gene sites was significantly enhanced in the knockout condition (Fig. 3L). Representative ATAC-seq track plots confirmed the specific increase in accessibility at the sites of key basal marker genes bound by POU2F3, including *Itga6* and *Trp63* (Supplementary Fig. 4J, K).

Together, these results demonstrate that POU2F3 represses basal lineage gene expression by directly binding to the regulatory elements and restricting chromatin accessibility of key basal genes.

### *Pou2f3* is Essential for Cell Type Distribution During Mammary Gland Development

Considering the essential function of *Pou2f3* in regulating luminal differentiation during mammary gland regeneration described above, it is paradoxical that it is not required for the physiological function of the organ. To resolve this paradox, we decided to perform a detailed phenotypic analysis of the *Pou2f3* null mammary glands during development. First, we examined *Pou2f3*^+/+^ and *Pou2f3*^+/-^ mice and did not observe any significant developmental differences between these two genotypes (Supplementary Fig. 5A-D). Therefore, we used heterozygous mice as controls for the null mice in the subsequent experiments. Carmine staining of the whole-mount mammary glands showed that epithelial morphogenesis was largely normal in *Pou2f3*^-/-^ mice when compared to the control mice, as ductal elongation and branch formation were both similar between them (Fig. 4A-D).

**Figure 4:**
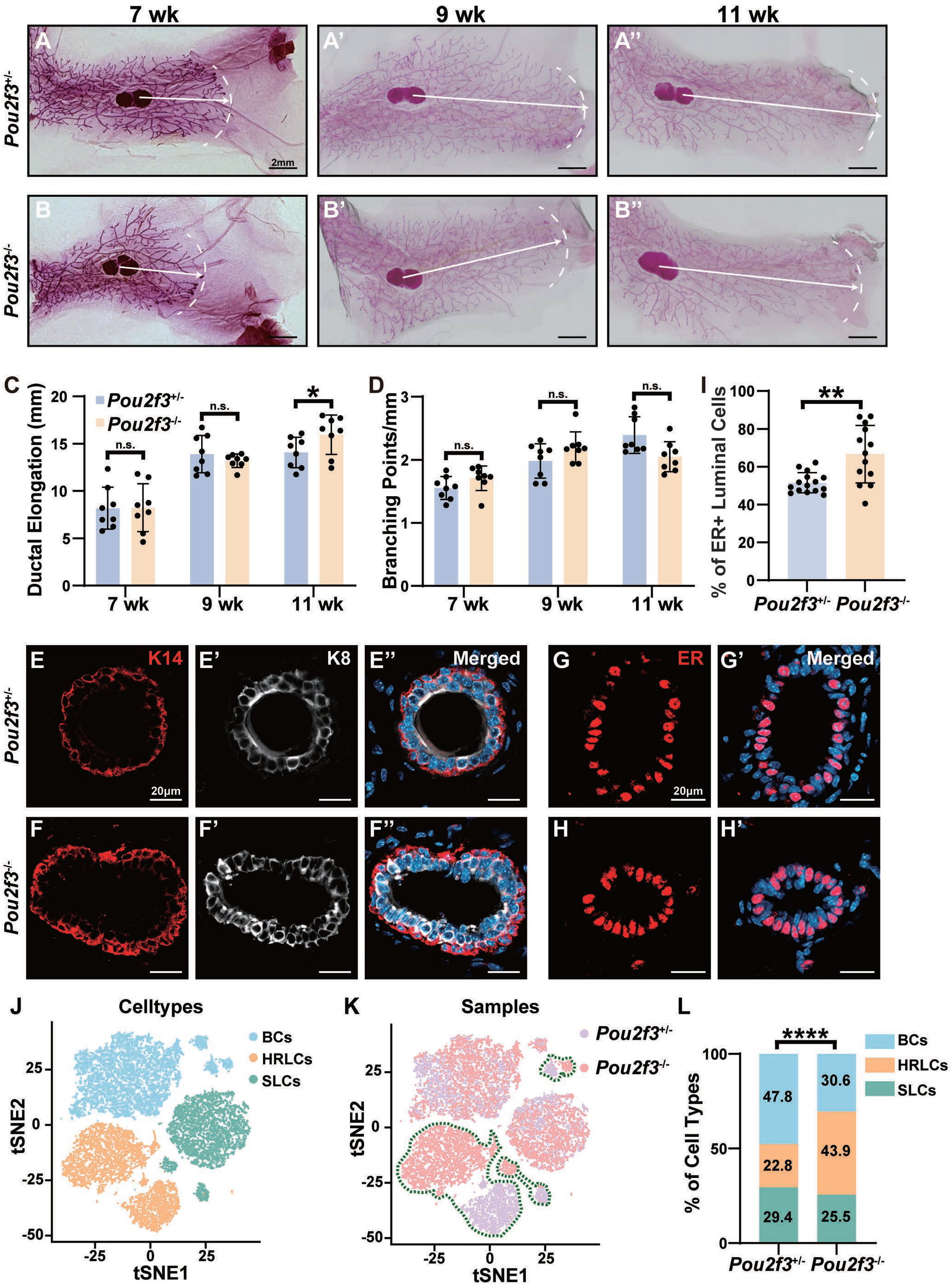
*Pou2f3* is Essential for Cell Type Distribution During Mammary Gland Development. (**A-D**) Mammary gland development in *Pou2f3* knockout and control mice. (**A, B**) Whole-mount carmine staining of mammary glands showing comparable branching morphology between control (**A**) and knockout (**B**) mice. (**C, D**) Quantification of ductal elongation (**C**) and branching points (**D**). Scale bars: 2 mm. N ≥ 3 mice. (**E-I**) Immunofluorescence staining of sections of the mammary epithelium using the indicated basal (**E**, **F**) and luminal (**E’**, **F’**, **G-H’**) cell markers from *Pou2f3* control (**E, G**) and knockout (**F, H**) mice. (**I**) Quantification of the percentage of ER+ luminal cells in the mammary epithelium based on FACS analysis. Scale bars: 20 µm. N ≥ 3 mice. (**J-L**) Single-cell RNA sequencing (scRNA-seq) analysis of mammary glands from *Pou2f3* knockout and control mice. (**J, K**) tSNE plots showing the distribution of cell types (**J**) and samples (**K**) of single cells from *Pou2f3* KO and control mice. The three circles with green dashed-lines highlight the non-overlapping, distinct clusters of cells that were supposed to be the same subtypes. (**L**) Percentage distributions of different cell subtypes in the mammary epithelium. Data are mean ± SD. Statistical analysis was performed using unpaired Student’s t-test. *p < 0.05; **p < 0.01; ***p < 0.001; ****p < 0.0001; n.s., not significant.

To determine whether cell differentiation was normal in mammary glands lacking *Pou2f3*, we used immunofluorescence confocal microscopy and examined the expression of a series of basal, luminal, and epithelial tissue polarity markers, including K14, K8, K18, ER, ECAD, and OCLN (Fig. 4E-H’, Supplementary Fig. 5E-J’). As expected, all these markers were expressed in the right cells and at the right locations, indicating that both basal and luminal cells were present and that they organized into a tissue architecture similarly in the *Pou2f3* null glands as in the control (Fig. 4E-H’, Supplementary Fig. 5E-J’). Curiously, we found that the percentage of ER^+^ luminal cells was increased by approximately 30% in the *Pou2f3* null glands compared to controls (Fig. 4G-I). When the percentage of luminal cells was quantified, we found it was also significantly increased in the *Pou2f3* null glands when compared to the control (Fig. 4E-F’’, Supplementary Fig. 5K). Using flow cytometry, we confirmed a small, though significant, increase in the percentage of luminal cells in the null glands (Supplementary Fig. 5L-N).

To further examine this phenotype, we measured the cell density of both basal and luminal epithelia in the 7-week mammary glands. We found that while basal cell density was similar in both genotypes, luminal cell density was significantly higher in the *Pou2f3* null gland than the control gland (Supplementary Fig. 5O). Together, these data show that the increase in ER^+^ cell percentage is due to an increase of luminal cells, rather than a decrease of basal cells, in the mutant mammary gland epithelium.

To validate the defect in cell type distribution and perform a detailed molecular analysis, we performed scRNA-seq on mammary epithelial cells harvested from *Pou2f3* control and null glands at the 7-week stage. Based on the expression features of specific marker genes, we classified the mammary epithelial cells into basal cells, HRLCs, and SLCs (Fig. 4J, K). Consistent with our immunofluorescence and FACS data, cell proportion analysis showed a significant increase in the percentage of luminal cells in *Pou2f3* knockout mice (Fig. 4L), with the increase in HRLCs being particularly pronounced.

Together, our results show that while *Pou2f3* is dispensable for epithelial morphogenesis, its absence alters cellular composition, molecular profiles, and potentially function of the main epithelial subtypes of cells. Furthermore, while *Pou2f3* loss leads to a reduction in luminal differentiation in regeneration, it appears to cause an opposite phenotype during development, leading to an increase of luminal percentage in the mutant epithelium.

### A POU2F3-TNF Feedback Loop Maintains Mammary Epithelial Homeostasis

To determine whether the abnormality in cell type proportions arises from a defective cell fate decision during the early differentiation stage or from a homeostatic imbalance occurring at a later stage, we used immunofluorescence and FACS analysis to examine the percentage of basal and luminal cells in the epithelium at the two-week stage, just when cell differentiation was completed but before epithelial branching began^35^. We found that the proportions of basal and luminal cells were similar in *Pou2f3* null and control epithelium (Supplementary Fig. 6A-F). Consistent with these results, we did not see any difference in basal or luminal cell density between control and null epithelia at the 2-week stage (Supplementary Fig. 6G). Together, these findings suggest that the change occurred during later stages, consistent with a defect in homeostatic maintenance.

We investigated whether the luminal cell increase in the *Pou2f3* null glands was due to increased proliferation or reduced apoptosis. However, a GSEA analysis of proliferation-related pathways, using the scRNA-seq data, were greatly reduced in the null luminal cells when compared to control (Fig. 5A). Consistent with these data, cell cycle analysis showed that the percentage of *Pou2f3* null luminal cells in the cell cycle (G2/M and S phases) was lower than control (Supplementary Fig. 6H). Likewise, Ki67 staining showed that cell proliferation was lower in *Pou2f3* null luminal cells than in control cells (Supplementary Fig. 6I). Moreover, TUNEL analysis showed that programmed cell death was higher in the mutant luminal cells, although in both genotypes of tissue, the percentage of cells undergoing cell death was overall extremely small at this stage (Supplementary Fig. 6J). Consistent with these results, colony-forming assays showed that *Pou2f3* null luminal cells formed a significantly fewer number of colonies than control luminal cells (Supplementary Fig. 6K-M). Together, these results show that the percentage increase in luminal cells is unlikely to result from changes in the luminal cells themselves.

**Figure 5:**
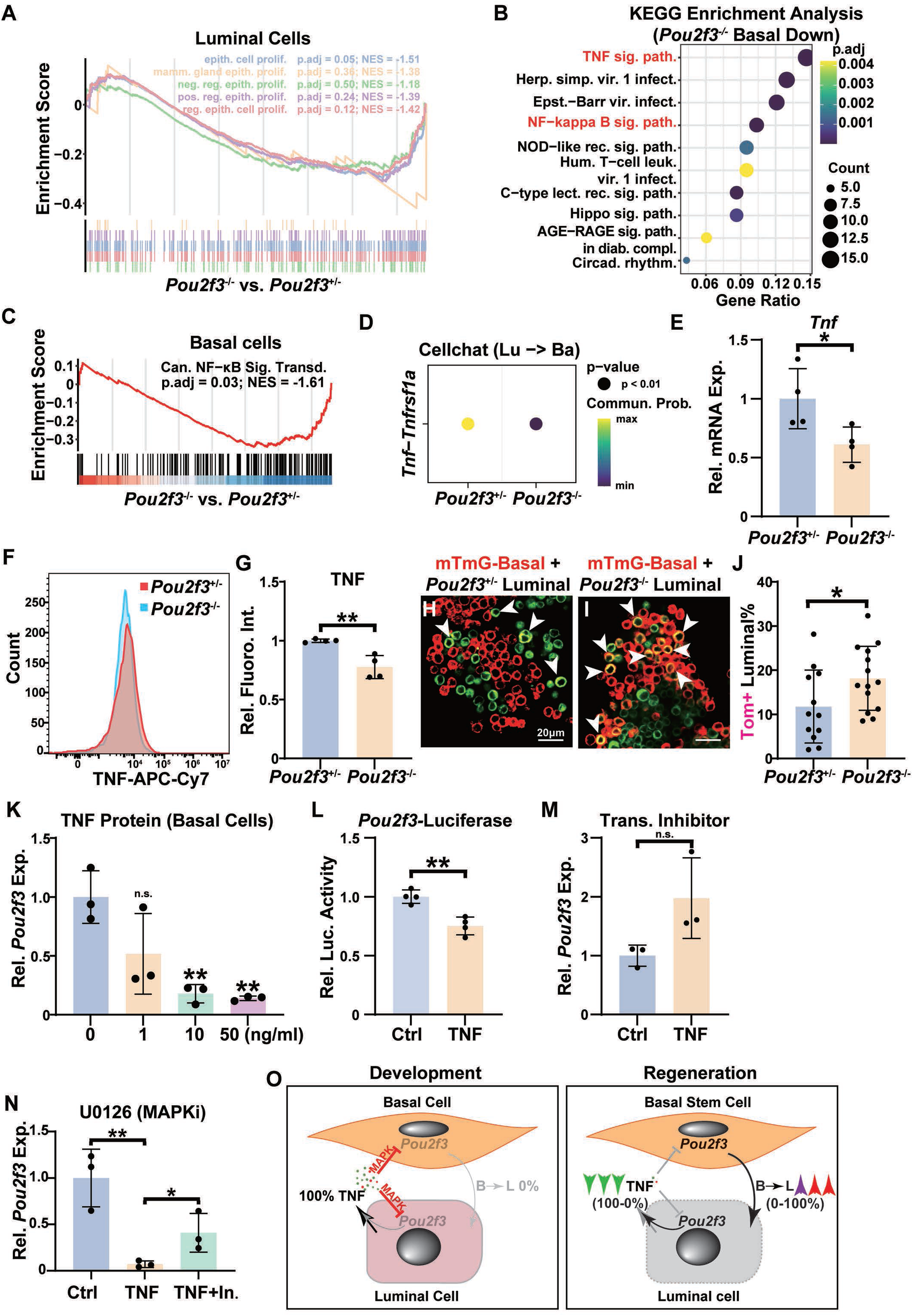
A POU2F3-TNF Feedback Loop Maintains Mammary Epithelial Homeostasis. (**A**) GSEA analysis of the proliferation-related pathways in luminal cells of the *Pou2f3* mutant mammary gland when compared with control mammary glands. (**B**) KEGG analysis of the downregulated genes in *Pou2f3* null basal cells, emphasizing the TNF signaling pathway and its downstream NFκB signaling pathway. (**C**) GSEA analysis of the NFκB signaling pathway genes in basal cells of the *Pou2f3* mutant mammary gland when compared with control mammary glands. (**D**) Cell-cell interaction analysis using CellChat, revealing reduced TNF signaling from luminal-to-basal cells in *Pou2f3* knockout mice compared to control. (**E**) Relative *Tnf* mRNA expression as revealed by qPCR reaction in luminal cells from *Pou2f3* knockout and control mice. N = 4. (**F**, **G**) TNF protein expression based on FACS analysis in luminal cells of the *Pou2f3* control and mutant mammary glands (**F**) and quantification of TNF fluorescent intensity (**G**). N = 4. (**H-J**) Co-culture of wild-type basal cells (red, derived from the mTmG mammary epithelium) with luminal cells derived from either *Pou2f3* control (**H**) or mutant (**I**) mammary epithelium. Basal and luminal cells were mixed at a 1:1 ratio and cultured. Luminal cells were detected using K8 immunofluorescence (green) at the end of the experiment. (**J**) Quantification of the percentage of luminal cells that were derived from basal cells. Note arrowheads denote select basal-derived luminal cells (both red- and green-colored). N ≥ 12. (**K**) Relative *Pou2f3* mRNA expression, as detected by qPCR, in basal cells cultured in medium with the indicated TNF protein concentration. N = 3. (**L**) Relative luciferase activity in HC11 cells carrying a construct expressing a luciferase under the control of the *Pou2f3* promoter, following TNF stimulation. N = 4. (**M, N**) Relative *Pou2f3* mRNA expression, as detected by qPCR, in HC11 cells cultured in medium with or without TNF protein and the indicated inhibitor. (**M**) Effect of the transcription inhibitor actinomycin D. (**N**) Effect of the MAPK inhibitors U0126. N = 3. (**O**) Model diagram of the mutual regulation of POU2F3 and TNF during mammary gland development and regeneration. Data are mean ± SD. Statistical analysis was performed using unpaired Student’s t-test. *p < 0.05; **p < 0.01; ***p < 0.001; ****p < 0.0001; n.s., not significant.

Therefore, we shifted our focus to basal cells. Using KEGG and GO enrichment analysis based on our scRNA-seq data, we found that downregulated genes were significantly enriched in the TNF and its downstream NFκB signaling pathways (Fig. 5B, Supplementary Fig. 7A). This was confirmed by GSEA analysis showing that the NFκB pathway was significantly reduced in *Pou2f3*^-/-^ basal cells (Fig. 5C). Considering the well-established role of TNF in inhibiting basal-to-luminal differentiation ^22^, we suspected that a reduction in TNF signaling may lead to a reduction in the inhibition of basal-to-luminal differentiation. This could reduce inhibition of basal-to-luminal conversion, resulting in a decreased basal fraction and an increased luminal fraction in the epithelium.

To test this possibility, we analyzed our scRNA-seq data and found that luminal-to-basal TNF-receptor interaction strength, based on the CellChat tool ^36^, was reduced in the mutant gland (Fig. 5D). This explains the observed reduction of TNF signaling in basal cells (Fig. 5B, Supplementary Fig. 7A) and predicts that TNF, normally expressed by luminal cells ^22^, is reduced in *Pou2f3* null glands. We tested this by examining the scRNA-seq data and found that *Tnf* mRNA expression is reduced in *Pou2f3* null luminal cells (Supplementary Fig. 7B). This was confirmed by qPCR reactions (Fig. 5E). Moreover, using FACS analysis, we found a 20% drop in TNF protein expression by *Pou2f3* null luminal cells (Fig. 5F, G). Thus, our results indicate that *Pou2f3* promotes *Tnf* mRNA and protein expression. Consistent with these results, *Tnf* expression is relatively highly expressed in the IMSCs where *Pou2f3* is upregulated during regeneration, especially when compared to basal cells where it is lowly expressed in basal cells during both development and regeneration (Supplementary Fig. 7C, D). Interestingly, the regulatory region of *Tnf* was absent from the POU2F3 binding sites based on our CUT&Tag experiment, suggesting that POU2F3’s promotion of TNF expression is likely an indirect consequence.

To directly determine whether TNF reduction due to *Pou2f3* loss in luminal cells would promote basal-to-luminal conversion, we lineage-traced basal cells, using them from *R26R*^mTmG^ female mouse mammary glands, and co-cultured them with luminal cells with or without *Pou2f3* function. As predicted, we found that basal-to-luminal conversion increased by around 67% when basal cells were co-cultured with *Pou2f3* null luminal cells when compared with control luminal cells (Fig. 5H-J).

At present, the basis of the well-known function of TNF inhibition of basal cell multipotency remains unknown. Given the essential role of *Pou2f3* in driving basal-to-luminal conversion, a tantalizing question is whether the well-known function of TNF inhibition of basal cell multipotency is mediated by blocking *Pou2f3* expression. If so, we would predict TNF signaling is reduced when basal cells are transitioning into IMSCs during mammary gland regeneration. This is indeed the case, as our KEGG analysis of downregulated pathways in IMSCs when compared to basal cells emphasized the TNF signaling pathway (Supplementary Fig. 7E). To directly test whether TNF inhibits *Pou2f3* expression, we added TNF protein to the culture medium of primary basal cells. We found that TNF inhibited *Pou2f3* mRNA expression in a dose-dependent manner (Fig. 5K). Similar inhibitory effects of TNF were also observed when the HC11 mammary cell line, which is more manipulatable than primary cells, was used (Supplementary Fig. 7F). Using luciferase reporter assays and transcriptional inhibition experiments, we confirmed that TNF represses *Pou2f3* transcription rather than regulating its mRNA degradation (Fig. 5L, M). To examine which of the downstream branches of TNF signaling (e.g., MAPK, NFκB, and JNK) were responsible for TNF inhibition of *Pou2f3* expression, we added their respective inhibitors to the culture medium of HC11 cells. We found that the addition of only U0126, a MAPK inhibitor, showed de-repression of *Pou2f3* mRNA expression (Fig. 5N, Supplementary Fig. 7G, H). The data thus suggest that TNF inhibits *Pou2f3* mRNA primarily through the MAPK signaling pathway.

In summary, we found that the abnormality in cell type proportions arises from an epithelial cell type imbalance occurred during homeostasis. More importantly, we identified a negative feedback loop where POU2F3 in luminal cells promotes TNF expression, which in turn signals back to basal cells to suppress *Pou2f3* expression and inhibit basal-to-luminal conversion, thereby maintaining homeostasis (Fig. 5O; see Discussion).

### Distinct Molecular Programs Govern Cell Fate in Organ Development and Regeneration

The above discovery that *Pou2f3* plays a critical role in regeneration but is dispensable for mammary gland development challenges the conventional wisdom that the two processes are mechanistically similar. Thus, we predicted that embryonic MSCs go through a different intermediate state to form luminal cells than that during regeneration, and this process is governed by a distinct differentiation program at the molecular level. To test this prediction, we first attempted to identify the intermediate state during luminal differentiation in development. We integrated published scRNA-seq datasets from embryonic day 16 (E16), embryonic day 18 (E18), postnatal day 4 (P4), and adult stages (Supplementary Fig. 8A) ^37^. The single cells were then scored based on their expression of basal and luminal marker genes (Supplementary Fig. 8B, C). Based on the scores, we partitioned them into embryonic (Embr), P4 basal (P4Ba) and luminal (P4Lu), and adult basal (AdBa) and luminal (AdLu) cell populations (Fig. 6A). Pseudotime analysis showed that cell fate had already diverged significantly by the P4 stage, forming independent lineage trajectories that lead to either adult basal or luminal cells (Fig. 6B). Cytotrace analysis revealed that the luminal lineage exhibited a progressively increasing differentiated state over developmental time (Fig. 6C, Supplementary Fig. 8D), indicating that P4 luminal cells are a key intermediate transitional state in the maturation of luminal cells.

**Figure 6:**
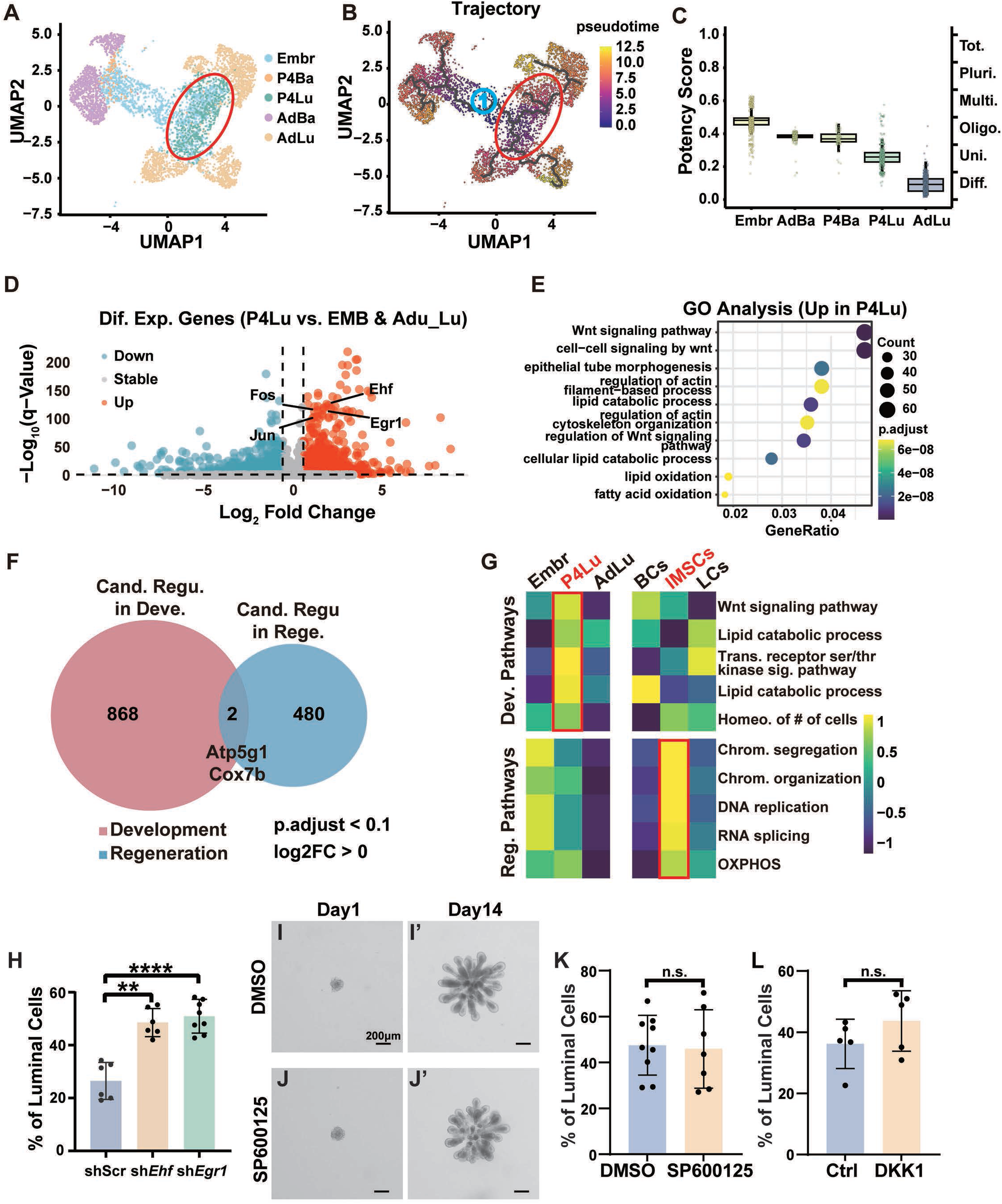
Distinct Molecular Programs Govern Cell Fate in Organ Development and Regeneration. (**A**) Single-cell RNA sequencing (scRNA-seq) data from embryonic day 16 (E16), embryonic day 18 (E18), postnatal day 4 (P4), and adult mammary glands were integrated and analyzed. Cells were classified into embryonic (Embr), P4 basal (P4Ba) and luminal cells (P4Lu), and adult basal (AdBa) and luminal cells (AdLu) based on transcriptomic signatures. (**B**) Pseudotime trajectory analysis revealed that the basal and luminal lineages diverge as early as P4, with cells gradually differentiating toward their respective adult identities. (**C**) Cytotrace analysis was used to assess cellular differentiation status. The results confirmed a progressive loss of plasticity from the embryonic to adult luminal state, identifying P4 luminal cells as an intermediate stage of luminal differentiation. (**D, E**) Differential gene expression analysis identified P4 luminal cell-specific markers (**D**). Gene enrichment analysis showed that these markers were primarily associated with Wnt signaling and oxidative stress pathways (**E**). (**F**) Venn diagrams illustrating the methods by which differentially expressed candidate regulators of development and regeneration were extracted. (**G**) Gene enrichment analysis demonstrated distinct signaling pathway activity: luminal cells in development were predominantly regulated by Wnt signaling, whereas luminal cells in regeneration were primarily associated with chromatin remodeling. (**H-L**) In vitro minigland regeneration assay where aggregates of basal cell were transfected with lentivirus expressing a scramble control, *Ehf*, or *Egr1* shRNA (**H**) or cultured in medium containing DMSO (**I, I’**) or the JNK pathway inhibitor SP600125 (**J, J’**). (**H, K, L**) Quantification of the percentage of luminal cells where *Ehf* or *Egr1* expressed was reduced using shRNA (**H**), where JNK inhibitor was used (**K**) or the WNT signaling inhibitor DKK1 (**L**) was used. N ≥ 5. Data are mean ± SD. Statistical analysis was performed using unpaired Student’s t-test. *p < 0.05; **p < 0.01; ***p < 0.001; ****p < 0.0001; n.s., not significant.

We then performed differential gene expression analysis on P4 luminal cells compared to embryonic and adult luminal cells. Consistent with our prediction that P4Lu-specific genes are candidate regulators of luminal differentiation, we found that *Ehf* and the JNK signaling component gene *Egr1*, which are known to be important for luminal cell differentiation ^38,39^, were significantly highly expressed in P4Lu cells (Fig. 6D, Supplementary Fig. 8E-F). Gene enrichment analysis showed that the most up- or down-regulated genes in P4Lu cells were vastly different from those during the regenerative process (Fig. 6E, Supplementary Fig. 8G; compared to Fig. 2B, Supplementary Fig. 2B). An analysis of the potential regulatory genes during development and regeneration showed only a minimal overlap between the two (Supplementary Fig. 8H, Fig. 6F). Using gene enrichment assay, we found that the signaling pathways represented by the marker genes of the two processes were highly specific, with development primarily involving signaling pathways such as WNT signaling, and regeneration primarily involving chromatin remodeling (Fig. 6G).

Next, we performed functional experiments to test the role of *Ehf* and *Egr1* in the regenerative process. Using qPCR, we confirmed the shRNAs for both genes effectively reduced their mRNA expression (Supplementary Fig. 9A, B). Interestingly, sh*Ehf* and sh*Egr1* organoids formed sparser though longer branches than normal by day 14 (Supplementary Fig. 9C-E’). As predicted, however, luminal differentiation in *Ehf* or *Egr1* knockdown organoids was not blocked; in fact, these organoids had a higher-than-normal luminal percentage (Supplementary Fig. 9F-H, Fig.6H). Thus, consistent with the branching phenotype observed in *Pou2f3*-GOF organoids (Supplementary Fig. 4C-D’’’), the observed anomaly in epithelial branching of sh*Ehf* and sh*Egr1* organoids is correlated with an overabundance of luminal cells in the organoid epithelium. These data thus confirmed that *Ehf* or *Egr1* function is not required for luminal differentiation in regeneration.

Moreover, we cultured basal cell organoids in a minigland regeneration assay in the presence of either a control (DMSO) or the JNK pathway inhibitor SP600125 (Fig. 6I-J). Interestingly, we found that SP600125 had no significant effect on the branching morphology of the organoids and did not inhibit luminal differentiation (Fig. 6K). Likewise, we tested the effect of the Wnt pathway, which was significantly enriched during development, using two different Wnt signaling pathway inhibitors, DKK1 and IWR-1, in the minigland assay. We found that inhibition of the Wnt pathway had no significant effect on the regenerative differentiation process (Supplementary Fig. 9I-Q, Fig. 6L).

We concluded that while both mammary gland development and regeneration proceed through an intermediate cell state, they are driven by distinct transcriptional programs. Specifically, developmental differentiation is regulated by Wnt signaling, and other pathways essential for cell-cell communication, while regeneration is mediated by chromatin remodeling driven by *Pou2f3*, that emphasizes fate change for rapid injury repair.

### Injury Repair, but not Development, of the Prostate and Pancreas Depends on *Pou2f3* function

We wondered whether the observation that developmental and regeneration programs are distinct, especially with their differential requirement for *Pou2f3*, is unique to the mammary gland or a general phenomenon for other vertebrate organs. We predicted that an organ lacking *Pou2f3* would develop relatively normally but would be defective in regeneration. We chose the prostate and pancreas, both of which are glandular organs with established injury and repair models ^40–42^, to test this prediction. As expected, we found that luminal differentiation was normal in the prostate lacking *Pou2f3*, as evidenced by FACS analysis and immunofluorescence, with normal epithelial architecture and cell composition (Fig. 7A, B, C, F, Supplementary Fig. 10A-B’’’, E). However, fewer luminal cells were replenished in the *Pou2f3*-null prostate compared to controls (Fig. 7D, E, F, Supplementary Fig. 10C-D’’’, E). Likewise, differentiation of AMY2A+ acinar cells was normal in the pancreas lacking *Pou2f3* (Fig. 7G-I’, L, Supplementary Fig. 10F-G’), but we found that the replenishment of damaged cells was significantly less in *Pou2f3* null than control pancreas after injury (Fig. 7J-N).

**Figure 7:**
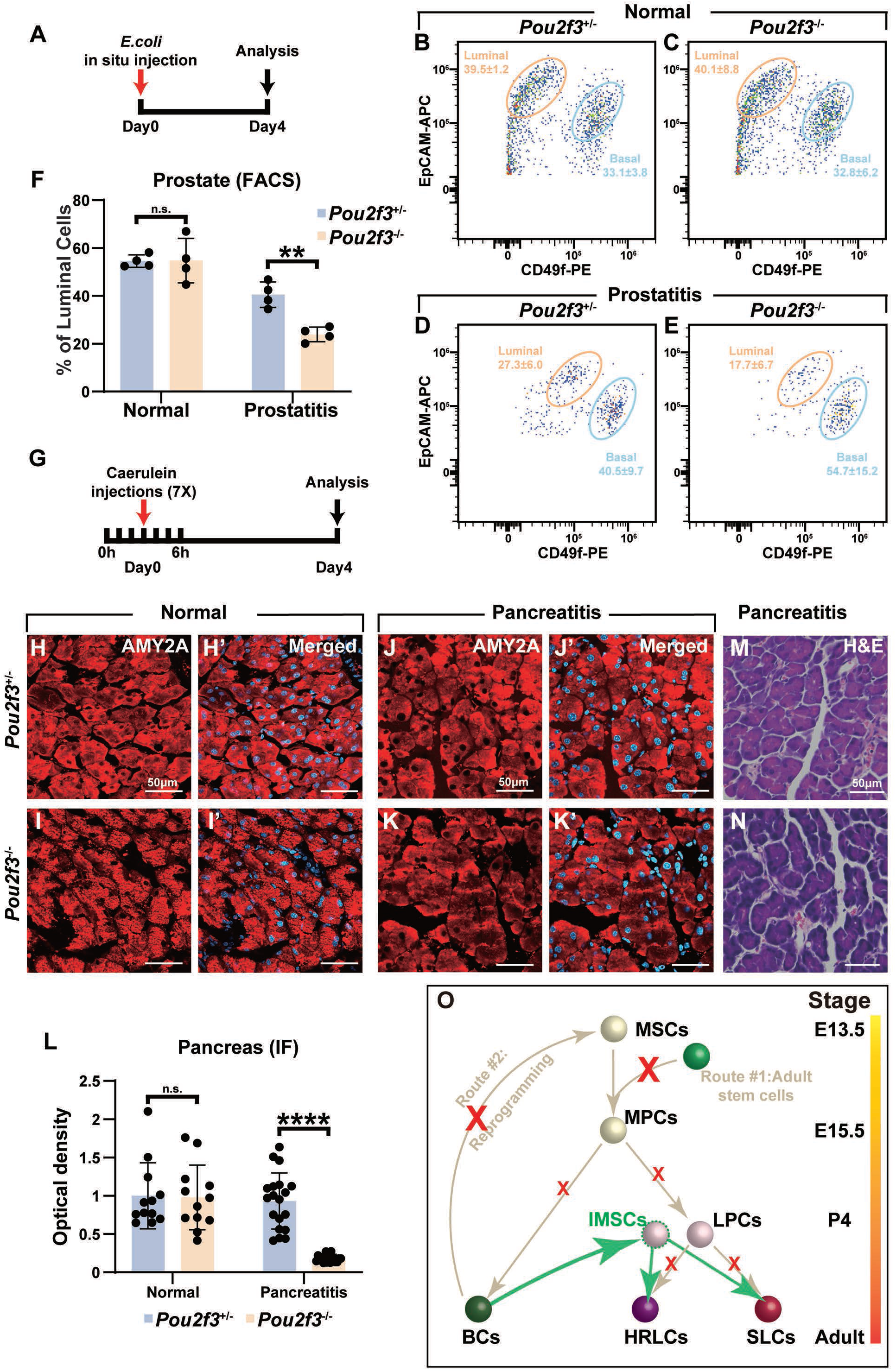
Injury Repair, but not Development, of the Prostate and Pancreas Depends on *Pou2f3* function. (**A**) Protocol diagram for prostatitis induction. (**B-F**) Assessment of the percentage of luminal cells, based on FACS analysis (**B-E**) in the *Pou2f3* control and null prostate under normal (**B, C**) or a prostatitis (**E, E**) condition. (**F**) Quantification of the percentage of luminal cells based on FACS. N ≥ 4 mice. (**G**) Protocol diagram for pancreatitis induction. (**H-L**) The percentage of AMY2A^+^ cells, based immunofluorescent microscopy (**H-K’**) in the *Pou2f3* control and null pancreas under normal (**H-I’**) or a pancreatitis (**J-K’**) condition. (**L**) Quantification of the percentage of luminal cells based on immunofluorescent microscopy. N ≥ 4 mice. (**M, N**) H&E staining of a paraffin section of the *Pou2f3* control and null pancreas after recovering from a pancreatitis condition. N ≥ 4 mice. (**O**) Diagram of a revised model of mammary gland stem cell differentiation during development and regeneration. Abbreviations: MSCs, mammary stem cells; MPCs, mammary progenitor cells; BCs, basal cells; HRLCs, hormone-responsive luminal cells; SLCs, secretory luminal cells. Data are mean ± SD. Statistical analysis was performed using unpaired Student’s t-test. **p < 0.01; ****p < 0.0001; n.s., not significant.

Together, the data show that distinct developmental and regeneration programs, which is independent and dependent, respectively, of *Pou2f3*, is a general phenomenon also observed in the prostate and pancreas. This provides a new model of how distinct biological processes can achieve similar outcomes through different molecular mechanisms (Fig. 7O) and suggests that this dichotomy is a general phenomenon in vertebrate organs.

## DISCUSSION

In this study, we addressed a central question in biology: are the molecular programs that govern embryonic development and adult tissue regeneration mechanistically similar? We demonstrate that despite a shared outcome of forming a mature epithelial structure, the mouse mammary gland utilizes distinct molecular programs for its development and regeneration. We identified a transient intermediate cell state that is essential for basal-to-luminal differentiation during regeneration and discovered that the transcription factor *Pou2f3* is a master regulator of this process. Surprisingly, we found that *Pou2f3* is largely dispensable for mammary gland development. We further revealed that POU2F3 functions as a transcriptional repressor that directly remodels chromatin, and is a key component of a novel feedback loop that maintains epithelial homeostasis. Finally, we showed that this dichotomy between developmental and regenerative programs is a general phenomenon that also applies to the prostate and pancreas. Our findings challenge the long-standing dogma that regeneration is simply a recapitulation of development, establishing a new paradigm that has broad implications for our understanding of tissue plasticity and disease (Fig. 7O).

### Distinct Cell State Transitions During Mammary Gland Development and Regeneration

We show that the course of cell fate or state changes during mammary gland regeneration is distinct from that during development. Specifically, during development, mammary stem cells (MSCs) progress through the stages of mammary progenitor cells (MPCs) to branch out, giving rise to basal cells (BCs) and luminal progenitors (LPs), the latter of which eventually differentiate into HRLCs and SLCs. By contrast, during regeneration, basal cells transition into IMSCs before becoming HRLCs and SLCs, without first becoming MSCs, MPCs, or LPCs, judging by various signatures including potency scores, trajectories, and molecular signatures (Fig. 7O).

Thus, our data challenge the current two schools of thought concerning the cell of origin of mammary gland regeneration, which argue that it is either adult stem cells or derivatives of reprogrammed basal cells ^18–20^. Moreover, despite their proposals of the origin of stem cells, both schools support the belief that stem cells follow a similar differentiation program in regeneration to the one in development ^21^. By contrast, our work shows that basal cells are directly converted into luminal cells in a process whereby POU2F3 restricts chromatin access to the basal programs to shut down its expression (Fig. 7O).

Based on scRNA-seq and developmental trajectory analysis, including pseudotime and RNA Velocity, we identified two distinct groups of transitional state cells, the IMSCs and P4Lu, that give rise to HRLCs and SLCs during mammary gland regeneration and development, respectively. Interestingly, the IMSCs are distinct from the “hybrid cells” previously identified as CD29^high^EpCAM^high^ based on FACS and proposed to be intermediate cells during basal-to-luminal conversion using the cytotoxin-mediated luminal-ablation model ^22^. One likely explanation for the distinction between the IMSCs and the hybrid cells is the method by which they were discovered -whereas the former was discovered by an unbiased approach using UMAP analysis based on the scRNA-seq data, the latter was based on FACS, despite subsequent gene discovery analysis based on scRNA-seq ^22^. This explains why IMSCs are unique from other cell subtypes, including mature or transitional basal and luminal cells, whereas hybrid cells fall into many different cell subtypes, including mature basal and luminal cells.

### The Molecular Function of POU2F3 in Basal-to-Luminal Cell Fate Decision

Our work provides a detailed molecular and cellular mechanism for how a single transcription factor, POU2F3, can drive a lineage switch during regeneration. *Pou2f3* is uniquely expressed in the IMSCs and its loss severely impairs luminal cell differentiation, leading to a failure of mammary gland regeneration. RNA-sequencing showed that *Pou2f3* promotes expression of genes in the luminal program. However, CUT&Tag analysis revealed that POU2F3 does not bind to the regulatory regions of most of these genes, suggesting the transcriptional regulation is secondary.

Instead, our data showed that POU2F3 binds predominantly to the distal regulatory regions of basal-specific genes. This binding leads to the active repression of the basal transcriptional program, as evidenced by our ATAC-seq data, which showed that the chromatin at these sites becomes significantly more accessible in the absence of POU2F3. These findings demonstrate that POU2F3’s primary function in this context is to silence the original basal cell identity, thereby enabling the cell to transition toward a new, luminal fate.

By contrast, during development luminal cells are derived from specified MSCs, rather than committed BCs. The basal program has yet to be initiated and, thus, does not require its shut down or POU2F3 function for the specified MSCs to take on the luminal lineage. The data also align with the known function of POU2F3 as a master regulator of lineage identity in other epithelial cells, such as chemosensory tuft cells, where its expression defines a specific cell fate ^29,30^. Together, our findings provide a clear example of how a transcription factor directly remodels chromatin to drive a cellular lineage switch.

### The POU2F3-TNF Feedback Loop and Epithelial Homeostasis

Our unexpected finding that *Pou2f3* is essential for regeneration but seemingly dispensable for normal development prompted us to investigate its function in adult tissue homeostasis. We discovered that while the morphology of the *Pou2f3* null mammary gland is largely normal, it has a mild defect in cell type proportions, with an overabundance, rather than a reduction as in regeneration, of luminal cells and an increased cell density. This is a mild phenotype that is often missed by gross morphological examinations. We show that this imbalance is neither a defect in cell differentiation during development, nor a result of changes in luminal cell proliferation or apoptosis but is caused by an altered basal-to-luminal conversion rate driven by a previously unrecognized cell-cell feedback loop.

Our data reveal a homeostatic circuit in which POU2F3, expressed in both basal and luminal populations, indirectly promotes the expression of TNF by luminal cells (Fig. 5O; Supplementary Fig. 11A). TNF, in turn, acts on basal cells to inhibit *Pou2f3* expression via a MAPK signaling pathway. This negative feedback loop acts as a homeostatic rheostat. Under normal conditions, a stable luminal cell population produces sufficient TNF to suppress *Pou2f3* expression in basal cells, thereby restricting basal-to-luminal differentiation and maintaining a stable cell ratio. When the luminal population is under-represented, for example, following injury or when basal cells are transplanted alone, the loss of luminal cells would reduce TNF signaling, thereby activating *Pou2f3* expression in basal cells and triggering the regenerative conversion to restore the luminal population (Fig. 5O; Supplementary Fig. 11A).

Importantly, the integration of both TNF inhibition on POU2F3 and POU2F3 promotion on TNF components into this unique feedback loop allows a rapid and sensitive response to turn on and off basal-to-luminal conversion activity. Such that, the switch could be turned on quickly when the epithelial tissue senses an insult, as measured by a drop in TNF expression due to reduced luminal population; and also turned off quickly when luminal population is replenished, as TNF expression is restored quickly due not only to increased luminal population but also POU2F3 promotion (Supplementary Fig. 11A). Considering most epithelial organs endure constant environmental insults, a similar feedback loop, potentially involving POU2F3 may operate in other epithelial organs. This is supported by our studies on the prostate and pancreas, where *Pou2f3* loss inhibits injury repair.

Finally, thanks to evolution, redundancy is often a common feature to ensure the robustness of an essential biological system. This is likely the case for the homeostatic loop that we discovered in the mammary epithelium. Specifically, a *Pou2f3*-independent mechanism appears to operate in parallel to convert basal to luminal cells. This is evidenced by the partial restoration of the luminal population when basal cells lacking *Pou2f3* were transplanted alone into the cleared fat-pad, and by the initially paradoxical increase of the luminal population observed in the *Pou2f3* null gland (Supplementary Fig. 11A). We await future studies to interrogate the nature of this hitherto unknown molecule that is redundant to *Pou2f3*.

### Distinct Mechanisms Underlying Cell Fate Decisions During Vertebrate Organogenesis and Regeneration

The central finding that *Pou2f3* is essential for cell fate decisions in regeneration but not for mammary gland development directly challenges the long-held belief that regeneration is a recapitulation of development. We demonstrate that while both processes generate a mature mammary gland, they achieve this outcome through fundamentally distinct molecular mechanisms. We used a systematic approach to identify the key regulatory programs in both contexts. During development, we identified an intermediate cell state at the P4 stage that is enriched for signaling pathways, such as WNT and JNK, that rely on community-based, cell-cell communication to regulate cell fate. By contrast, the intermediate state during regeneration is driven by chromatin remodeling orchestrated by *Pou2f3*, an “on-demand” mechanism that can be activated quickly and robustly in response to tissue loss. This dichotomy is not a unique phenomenon to the mammary gland. Our functional studies in the prostate and pancreas confirmed that their glandular development is normal in the absence of *Pou2f3*, whereas their regenerative capacity is severely compromised.

Our work on understanding cell fate decisions are consistent with recent studies showing that organ morphogenesis may be modified to suit the changing contexts during regeneration ^2,42,43^. Together, these studies establish a new paradigm for vertebrate organogenesis: development and regeneration are not two sides of the same coin but are instead governed by distinct molecular programs. This finding is of broad significance as it provides a new conceptual framework for understanding the fundamental processes of organogenesis and tissue repair across different organ systems.

Moreover, our work provides critical insights that have profound implications for regenerative medicine, stem cell biology, and disease. For decades, researchers in regenerative medicine have pursued the reactivation of developmental programs to promote tissue repair. Our findings suggest that this approach may be misguided. Instead of trying to restart the complex, multi-layered programs of development, a more effective strategy for treating acute tissue loss may be to specifically target the unique, streamlined regenerative pathways that are designed for repair. Our identification of a key transcriptional repressor, POU2F3, and a novel homeostatic feedback loop, provides concrete therapeutic targets for modulating cell plasticity and restoring tissue balance.

Finally, this study contributes significantly to the ongoing debate about the identity and function of adult stem cells. Our data support the idea that adult epithelial plasticity is a specialized, context-dependent process that is mechanistically distinct from the lineage decisions of embryonic development. The subtle homeostatic defect we observe in the *Pou2f3* null mammary gland suggests that a compromised regenerative program, even if it doesn’t cause a major developmental failure, could be a root cause of chronic tissue dysfunction or predispose to diseases like cancer, where a loss of cell population control is a hallmark.

### Limitations of the Study

While our study provides mechanistic insight into how *Pou2f3* governs basal-to-luminal differentiation and reveals distinct transcriptional programs between development and regeneration, several limitations should be acknowledged. First, although the transplantation and organoid assays effectively model mammary gland regeneration, they may not fully recapitulate the physiological microenvironment of injury-induced regeneration in vivo. Second, our single-cell transcriptomic analyses identify an intermediate cell state and its candidate regulators, yet functional validation was focused primarily on *Pou2f3*; the roles of other transcription factors enriched in this state remain to be determined. Third, although CUT&Tag and ATAC-seq analyses indicate that POU2F3 represses basal gene programs by restricting chromatin accessibility, additional biochemical studies will be required to define its interacting partners and chromatin-remodeling complexes. Finally, while we observed similar context-specific requirements for *Pou2f3* in prostate and pancreas regeneration, whether this paradigm extends to non-glandular tissues remains to be tested. Addressing these limitations will further refine our understanding of how distinct molecular programs control cell fate across regenerative and developmental contexts.

### Concluding Remarks

Together, our findings establish a new conceptual framework in which organ regeneration is not merely a reiteration of developmental processes but a distinct, context-dependent program optimized for rapid restoration of tissue integrity. By identifying *Pou2f3* as a pivotal regulator that orchestrates chromatin remodeling and represses basal identity to enable luminal differentiation, we uncover a molecular logic that distinguishes regenerative plasticity from developmental lineage specification. This paradigm challenges long-standing assumptions in stem cell and developmental biology and provides a foundation for re-evaluating how cell identity is maintained or reprogrammed in adult tissues. More broadly, delineating the divergent mechanisms governing development and regeneration may inform strategies to restore tissue function after injury or degeneration, with implications for regenerative medicine, cancer biology, and epithelial homeostasis.

## MATERIALS AND METHODS

### Mouse Strains and Animal Husbandry

Mice carrying the *R26R*^mT/mG^ reporter allele (JAX Mice, Stock No. 007576) ^25^ were procured from the Jackson Laboratory. *Pou2f3* knock-out mice were purchased from Cyagen Biosciences. C57BL/6J and BALB/c-Nude mice were obtained from GemPharmatech. All mice were housed in a pathogen-free vivarium with controlled temperature (18–23 °C), humidity (40–60%), and a 12-hour light, 12-hour dark cycle. Genotyping was performed according to protocols provided by the respective suppliers, with primer information in the Supplementary Table 2. All procedures were approved by the South China University Institutional Animal Care and Use Committee (IACUC# 2024#67) and conformed to ethical guidelines for animal research.

### Lactation and Pup Body Weight Measurement

For lactation capability experiments, 8-week-old female *Pou2f3* heterozygous (*Pou2f3*^+/-^) and mutant (*Pou2f3*^-/-^) mice were mated with wild-type male mice. Litter size and pup survival were recorded on the day of delivery and at postnatal day 21 (weaning). Pup body weights were also measured at weaning.

### Preparation of Mammary Gland Organoids and Epithelial Cells

Primary mammary gland epithelial cells and organoids were isolated as previously described ^32^. Briefly, inguinal (#4) and thoracic (#3) mammary glands were collected from 8 to 10-week-old female mice and minced with a razor blade. The minced tissue was digested in 10 mL of digestion solution (DMEM/F12 supplemented with 2 mg/mL collagenase (Sigma, Cat# C5138), 2 mg/mL Trypsin (Gibco, Cat# 27250018), 5 μg/mL insulin (Yeasen, Cat# 40107ES25), 5% fetal bovine serum (FBS) (Gibco, Cat# 10099141), and 50 μg/mL Gentamicin) for 30 minutes at 37°C with gentle shaking. Following digestion, mammary organoids were pelleted by centrifugation at 450 × *g* for 10 minutes and purified by 3–5 rounds of differential centrifugation, with washes in cold FACS buffer. To obtain single cells for subsequent sorting, purified organoids were dissociated with 0.25% Trypsin/EDTA (Meilunbio, Cat# MA0233) for 12 minutes at 37°C.

Alternatively, mammary organoids were generated from reaggregates of flow-sorted single cells. A mixture of 500 basal cells (*R26R*^mT/mG^-positive) and 500 luminal cells (*Pou2f3*^+/-^ or *Pou2f3*^-/-^) was aggregated overnight at 37°C in 96-well ultra-low attachment plates (Jet Biofil, Cat# TCP130096) to form spheroids. Aggregates were then embedded in 50 µL of phenol red-free basement membrane matrix (Matrigel, NEST Biotechnology Cat# 211282) in an eight-well chamber slide. Cultures were maintained in phenol red-free DMEM/F-12 (Procell, Cat# PM150316) supplemented with 1% HEPES (Procell, Cat# PB180325), 1% L-Ala-Gln, 1% P/S, N2, and B27, along with 100 ng/mL Nrg1 (ABclonal, Cat# RP01825), 100 ng/mL Noggin (ABclonal, Cat# RP01308), and 100 ng/mL R-spondin 1. The medium was changed every three days. On day 7, organoids were released from the matrix by digestion with 2 U/mL Dispase (Corning, Cat# 354235).

### Mammary Gland Transplantation

For mammary gland regeneration transplantation, 3-week-old female nude mice with cleared fat pads were used as hosts. A total of 10,000 mammary gland epithelial cells were injected into the cleared fat pad, and the fat pads containing reconstituted mammary glands were harvested after a specified period. The collected mammary glands were placed on adhesive glass slides for fluorescence imaging or were subjected to whole-mount carmine staining. Carmine staining involved fixing the glands in Carnoy’s fixative solution, followed by rehydration, staining with carmine, and subsequent dehydration and degreasing prior to imaging. Imaging of both fluorescent and stained samples was performed using a Zeiss Axio Zoom V16 stereoscope

### In Vitro Mammary Basal Cell Differentiation Assay (Minigland Culture)

The minigland culture method was adapted from previously established protocols ^31^. L Wnt-3A cells were cultured for 7 days to collect the conditioned medium, which was used for the culture of mammary basal cells. Briefly, 3000 flow-sorted basal cells were resuspended in Mammary sphere formation medium and seeded into 96-well plates pre-coated with 30 μL of growth factor-reduced Matrigel. The culture was maintained for 7 days to allow mammary sphere formation. Subsequently, spheres were released from the Matrigel by incubation with 2 U/mL dispase at 37°C for 30 minutes. Single or multiple spheres were then transferred to a 24-well plate containing 40 μL ECM mixture (Matrigel:Collagen I, 3:1) and incubated in a 37°C metal bath to solidify the ECM mixture. After solidification, 500 μL of culture medium was added to each well. From days 0-6, the culture medium was replaced with branching medium and refreshed daily; from days 6-10, elongation medium was used and replaced every other day; and from days 10-13, pseudo-estrus cycle medium was applied with daily medium changes. Details of the culture medium compositions can be found in the Supplementary Table 3.

For lentiviral infection, lentivirus was mixed with basal cells and seeded into 96-well plates pre-coated with 30 μL of Matrigel. After 7 days, organoids expressing fluorescence were selected for subsequent experiments.

For minigland harvesting, the organoids were released from the ECM mixture using 2 U/mL dispase at 37°C for 30 minutes. The released miniglands were then dissociated into single cells by treatment with 0.25% Trypsin/EDTA for 2 minutes at 37°C for subsequent sorting.

### Cell Line Cultures

HEK293T cells (Procell, Cat# CL-0005) were cultured in DMEM (Gibco, Cat# C12430500BT), supplemented with 10% FBS (Shanghai Life-iLab Biotech Co., Ltd, Cat# AC03L155), 1% sodium pyruvate, 1% non-essential amino acid, 1% L-Ala-Gln, and 1% P/S. L Wnt-3A cells (Procell, Cat# CL-0375) were cultured in DMEM (Gibco, Cat# C12430500BT), supplemented with 10% FBS (Shanghai Life-iLab Biotech Co., Ltd, Cat# AC03L155), 1% sodium pyruvate, 1% non-essential amino acid, 1% L-Ala-Gln, 1% P/S and 0.4 mg/mL G418 (Meilunbio, Cat# MA0321). HC11 cells were cultured in RPMI 1640 (Gibco, Cat# C11875500CP), supplemented with 10% FBS, 10 ng/mL EGF, 5 μg/mL insulin, 1% L-Ala-Gln, and 1% P/S. All cell lines were routinely tested for mycoplasma contamination using PCR.

### Subcloning and Construct Production

shRNA sequences were designed based on the BLOCK-iT™ RNAi Designer platform (**Thermo Fisher Scientific**). The pLKO.1 vector (Addgene, Cat# 8453) was digested with AgeI (Abclonal, Cat# RK21125) and EcoRI (Abclonal, Cat# RK21102), then ligated with annealed shRNA fragments using T4 DNA ligase (Abclonal, Cat# RK21501). Correct ligation was validated by Sanger sequencing. Knockdown efficiency was evaluated in basal cells, and the construct with the highest efficiency was selected for subsequent experiments. For construction of overexpression vectors, the full-length coding sequence (CDS) of *Pou2f3* was cloned into the pLeGO-SFFV-P2A-mCherry vector using the ClonExpress Ultra One Step Cloning Kit V3 (Vazyme Biotech co., ltd., Cat# C117). Plasmids were prepared using an endotoxin-free plasmid extraction kit (Magen, Cat# P1112).

The sequences of shRNAs used were:

Scramble shRNA: 5’-CCTAAGGTTAAGTCGCCCTCG-3’.

*Pou2f3*-shRNA: 5’-GCAGAGACGCATTAAGCTAGG-3’.

*Ehf*-shRNA: 5’-GCAGCTGGGTTACTCCTGTGT-3’.

*Egr1*-shRNA: 5’-GCCTCGTGAGCATGACCAATC-3’.

### Lentivirus Production

HEK293T cells were co-transfected with pMD2.G (Addgene, Cat# 12259), psPAX2 (Addgene, Cat# 12260), and the transfer plasmids, using the Eztrans transfection reagent (Shanghai Life-iLab Biotech Co., Ltd, Cat# AC04L099). Viral supernatant was harvested twice at 60 hours and 88 hours post-transfection. The lentivirus was concentrated by ultracentrifugation at 27,000 rpm for 2 hours and resuspended in DMEM/F12 containing 10% FBS. Aliquots were stored at −80°C.

### Immunofluorescence Analysis

Tissue samples were harvested and fixed in 4% paraformaldehyde (PFA) overnight at 4°C. Tissues were then equilibrated in 30% sucrose prepared in PBS overnight and embedded in OCT compound (Servicebio, Cat# G6059) for cryosectioning. Ten μm thick frozen sections were prepared using a Leica cryostat. Sections were permeabilized with 0.5% Triton X-100 in PBS for 45 minutes and blocked for 2 hours at room temperature (RT) in PBS containing 10% goat serum and 0.2% Tween-20. Primary antibodies (Supplementary Table 4) were applied and incubated overnight at 4°C. Following primary antibody incubation, sections were incubated with secondary antibodies for 2 hours at RT. Nuclei were counterstained with DAPI.

For whole-mount immunofluorescence of organoids, a suspension staining protocol was used ^44^. Organoids were fixed with 4% PFA at 4°C for 45 minutes. Fixed organoids were then transferred to an ultra-low attachment 24-well plate and incubated sequentially with primary and secondary antibodies. Nuclei were counterstained with DAPI. For clearing, organoids were resuspended in a glycerol-fructose solution after removing the final wash buffer.

Confocal microscopy was performed using either a Zeiss LSM 900 Confocal or a Nikon CSU-W1 Sora Confocal system.

### Fluorescence-Activated Cell Sorting (FACS)

Dissociated single-cell suspensions from primary mammary glands, prostate tissues, and cultured miniglands were incubated with specific antibodies (Supplementary Table 3) on ice for 40 minutes. After incubation, cells were washed and resuspended in FACS buffer (PBS containing 2% FBS and 1% P/S) and filtered through a 40-μm mesh. Sorting was performed on a CytoFLEX SRT (Beckman Coulter), and data were analyzed using FlowJo 10 software.

### Quantitative Real-Time PCR (qPCR)

Total RNA was extracted from cells using the RNA extraction kit (ABclonal, Cat# RK30120) according to the manufacturer’s instructions. Equal amounts of RNA were reverse transcribed into cDNA using the ABScript Neo RT Master Mix for qPCR with gDNA Remover (ABclonal, Cat# RK20433). The resulting cDNAs were then used for qPCR reactions with the 2X Universal SYBR Green Fast qPCR Mix (ABclonal, Cat# RK21203). qPCR was performed on an Agilent AriaMx Real-Time System. Gene expression levels were normalized to *Hprt*. Primer sequences are provided in the Supplementary Table 5.

### Construction of cDNA Libraries and Bulk RNA Sequencing

Total RNA was isolated from miniglands using the MicroElute RNA extraction kit (ABclonal, Cat# RK30124). Equal quantities of RNA were then reverse transcribed into cDNA using the Low Input cDNA Synthesis & Amplification Module (ABclonal, Cat# RK20310). cDNA quantification was performed with the Equalbit dsDNA HS Assay Kit (Vazyme, Cat# EQ111-01). Subsequently, 1 ng of cDNA was used for library construction with the One-step DNA Library Prep Kit for Illumina V2 (ABclonal, Cat# RK20237). Indexed libraries were pooled and sequenced on the Illumina NovaSeq 6000 platform using a 2×150 paired-end (PE) configuration by GENEWIZ (Suzhou, China).

Raw RNA-seq reads were subjected to quality control using FastQC (v0.11.9), and adapter trimming was performed with Trim Galore! (v0.6.7). Clean reads were aligned to the mouse reference genome (GRCm39) using the Subread package (v2.0.3). Gene-level read counts were obtained using featureCounts with the GENCODE annotation file (vM30). Differential gene expression analysis was conducted using DESeq2 (v1.38.1), and Gene Set Enrichment Analysis (GSEA) was performed with the clusterProfiler R package (v4.6.2).

### Single-Cell RNA Sequencing and Bioinformatics Analysis

For single-cell RNA sequencing, the Cell Ranger tool (v7.1.0) was used for quality control and quantification of the raw sequencing data from 10x Genomics. The Seurat R package (v5) was utilized for data input, quality control, clustering, and differential gene expression analysis ^45^. Basal and luminal cell marker genes, 1814 and 2139 genes, respectively, in transplanted samples were identified by differential analysis based on a previously published adult mammary gland single-cell transcriptome dataset ^26^. For embryonic mammary gland development analysis, the GSE1111113 dataset was used, and the Canonical Correlation Analysis (CCA) method in Seurat was applied to integrate different samples.

Downstream analyses were performed using several R packages: Cytotrace (v0.1.0) was used to assess cell stemness scores ^27^; RNA velocity was inferred using velocyto ^28^; Monocle 3 (v1.2.9) was applied to infer cell differentiation trajectories ^46^; and SCENIC (v1.2.6) was employed for transcription factor activity inference ^47^. Gene Ontology (GO), Kyoto Encyclopedia of Genes and Genomes (KEGG), and GSEA analyses were performed using the clusterProfiler (v4.6.2) ^48^. Cell-cell communication analysis was conducted using the CellChat (v2.1.2) R package ^36^.

### CUT&Tag and ATAC-seq

For CUT&Tag experiments, *Pou2f3*-/- basal cells were transduced with a POU2F3-FLAG lentiviral vector. Following selection, 100,000 FLAG-positive cells were sorted per sample. CUT&Tag libraries were prepared using the NovoNGS® CUT&Tag 4.0 High-Sensitivity Kit for Illumina (Novoprotein, Cat# N259-YH01) according to the manufacturer’s protocol. The antibody used for immunoprecipitation is detailed in the Supplementary Table 2.

For ATAC-seq experiments, 1,000 cells were sorted from miniglands for each sample. Libraries were constructed using the ATAC-Seq Lib Prep Kit for Illumina (ABclonal, Cat# RK21509) following the manufacturer’s protocol.

Indexed libraries were pooled and sequenced on the Illumina NovaSeq 6000 platform using a 2×150 paired-end (PE) configuration by GENEWIZ. Raw CUT&Tag and ATAC-seq reads were first subjected to quality control with FastQC (v0.11.9), and sequencing adapters were trimmed using Trim Galore! (v0.6.7). Cleaned reads were aligned to the mouse reference genome (mm39) using Bowtie2 (v2.4.5) for CUT&Tag and BWA-MEM (v0.7.17) for ATAC-seq. Low-quality reads and PCR duplicates were filtered with SAMtools (v1.16.1). Peak calling was performed using MACS2 (v2.2.9). Identified peaks were functionally annotated with ChIPseeker (v1.36.0). Normalized bigWig tracks were generated with deepTools bamCoverage (v3.5.1) for downstream analyses. To evaluate chromatin accessibility around CUT&Tag-defined peaks, each peak region was symmetrically extended and overlapping ATAC-seq signal intensities from bigWig files were extracted using rtracklayer (v1.60.2) and processed with GenomicRanges (v1.52.1) in R. Signal values were summarized into per-base average accessibility scores.

### Growth Factor Stimulation

Basal cells were cultured as spheres. The spheres were then transferred to basal medium supplemented with 2% growth factor reduced Matrigel for 24 h and subsequently treated with 10 ng/mL TNF (ABclonal, Cat# RP01071) for 24 h prior to RNA extraction.

HC11 cells were starved in serum-free medium for 16 h, followed by stimulation with 10 ng/mL TNF for 24 h. For transcription assays, serum-starved cells were pretreated with 1 µg/mL actinomycin D (MedChemExpress, Cat# HY-17559) for 1 h, followed by 10 ng/mL TNF stimulation for 6 h. For signal transduction inhibition experiments, cells were treated with 10 ng/mL TNF in the presence or absence of the following inhibitors for 24 h: 30 µM JSH-23 (MedChemExpress, Cat# HY-13982), 1 µM SP600125 (MedChemExpress, Cat# HY-12041), or 10 µM U0126 (MedChemExpress, Cat# HY-12031).

### Dual-Luciferase Reporter Assay

The *Pou2f3* promoter was cloned into the pGL4.10 vector. HC11 cells were co-transfected with pGL4.10 and the control vector pGL4.74 using a liposomal transfection reagent (Yeasen, Cat# 40802ES03). Following serum starvation, cells were stimulated with 10 ng/mL TNF. Luciferase activity was measured using the Dual-Luciferase Reporter Assay Kit (Beyotime, Cat# RG027) according to the manufacturer’s protocol.

### Preparation of Prostate Epithelial Cells and Injury Model

Prostate tissues were harvested and incubated in 1 mL of collagenase/hyaluronidase solution (DMEM/F12, containing 1x collagenase/hyaluronidase (Stemcell, Cat# 07912), 10 µM Y27632 (Selleck, Cat# S1049), and 50 μg/mL Gentamicin) for 45 minutes at 37°C with gentle shaking. Prostate organoids were pelleted by centrifugation at 500 × *g* for 3 minutes and purified by differential centrifugation. Organoids were digested in 0.25% Trypsin/EDTA for 6 minutes at 37°C to obtain single cells for subsequent sorting.

Prostatitis was induced by injection of 25 μl of a 1 × 10^10^ cfu/ml E. coli (DH5α) suspension into the anterior prostate of the male mouse ^40^. Four days later, mice were anesthetized, and the surgical site was prepared by shaving and disinfecting the abdominal area before making a small incision to expose the prostate. After the injection, the incision was sutured, and the mice were placed on a warming plate and returned to their cages.

### Pancreas injury model

Pancreatitis was induced by administering seven hourly intraperitoneal injections of 100 μg/kg cerulein (Meilunbio, MB2573) to the mice ^41^. The mice were euthanized and their pancreases collected 4 days following the first cerulein injection. Mice were monitored daily for changes in body weight and for signs of distress, including diarrhea, piloerection, lethargy, and periorbital exudates.

### Quantification and Statistical Analysis

Sample size for each figure is denoted in the figure legends. Statistical significance between conditions was assessed by two-tailed Student’s *t*-tests. All error bars represent SD, and significance is denoted as \**P* < 0.05, \*\**P* < 0.01, \*\*\**P* < 0.001 and **** *P* < 0.0001. n.s. denotes not significant.

## Data Availability

The bulk RNA sequencing and single cell RNA sequencing data have been deposited in NCBI’s Gene Expression Omnibus (GEO) repository and are accessible through GEO Series accession numbers (Accession number: GSE309852). All data supporting the conclusions of this study are provided in the main text or supplementary materials.

## ACKNOWLEDGEMENTS

We extend our gratitude to the Lu lab members for their insightful discussions. Special thanks to the Molecular Imaging Core Facility and the Molecular and Cell Biology Core Facility at Hengyang Medical School, South China University, for technical assistance. This research was supported by grants from the National Science Foundation of China (32470886 to P.L. and 82330035 to K.X.) and a key grant from the National Science Foundation of Hunan Province (2025JJ30012 to P.L.).

## AUTHOR CONTRIBUTIONS

R.M., Acquisition of majority of the data, writing, and review.

Z.D., Assistance on bioinformatics analysis.

Y.T., Phenotypic characterization using the mini-gland model; Characterization of the injury and repair models in the prostate and pancreas.

J.C., Characterization of the lactational mammary glands.

J.Z., Phenotypic characterization using the mini-gland model.

D.F., scRNA-seq and bioinformatics analysis on basal-based transplants on day 5.

C.X., Characterization of basal-to-luminal kinetics after transplantation.

J.Z., X.Z., K.X., intellectual contributions, review.

P.L., Conceptualization, funding acquisition, data curation, and writing.

