## Supplemental Figure for "Distinct Programs Drive Organ Development and Regeneration"

**Supplementary Figure 1: Identification of a Novel Intermediate Cell State During Mammary Gland Regeneration.**

(**A, B**) Violin plots showing the expression levels of basal or luminal markers in basal, luminal, and intermediate state cells in the basal-and-luminal (BaLu) (**A**) and basal-alone (Ba) (**B**) samples. Abbreviations: BCs, basal cells; LCs, luminal cells; IMSCs, intermediate state cells.

(**C**) UMAP plots showing the expression of *Elf5* and *Esr1,* marking the HRLC and SLC populations, respectively, in single cells of the Ba samples.

(**D, E**) UMAP plots showing the cell cycle phases of the single cells from the BaLu (**D**) and Ba (**E**) samples.

(**F**, **G**) Scoring of single cells on the UMAP plot (**F**) and the epithelial subtypes (**G**) from the Ba samples using hybrid-state cell markers from a published study. The score was calculated using 239 marker genes selected from the hybrid cells (CD29$^{high}$EpCAM$^{high}$).

(**H-K**) UMAP plots showing the expression of previously identified mammary stem cell markers, including *Tspan8*, *Procr*, *Bcl11b*, and *Lgr5* in the cell clusters from the Ba samples.

**Supplementary Figure 2: The POU Domain Transcription Factor *Pou2f3* is a Key Regulator of the Intermediate State.**

(**A**) Heatmap analysis of top 25 most upregulated and 25 most downregulated genes in IMSCs when compared with basal cells and luminal cells.

(**B**) GO enrichment analysis of the downregulated genes in IMSCs.

(**C, D**) UMAP plots showing the expression of the indicated candidate transcription regulators in single cells from the BaLu (**C**) and Ba (**D**) samples.

**Supplementary Figure 3: POU2F3 Promotes Basal-to-Luminal Conversion by Restricting Chromatin Accessibility to the Basal Transcriptional Program.**

(**A**, **B**) *Pou2f3* mRNA expression in different cell types at the 7-week-stage during development (**A**) and during mammary gland regeneration (**B**) based on scRNA-sequencing at these stages.

(**C-E**) Generation and validation of *Pou2f3* knockout mice. (**C**) Design of the knockout construct. (**D**) Genotyping validation by PCR reactions of *Pou2f3* knockout in mouse pups. (**E**) Detection of *Pou2f3* mRNA expression by qPCR reactions in control and null mammary epithelial cells. N= 3.

(**F**, **G**) Assessment of the lactation capacity of *Pou2f3* knockout mice. Quantification of the survival rates (**F,** N=3) and the weight (**G,** N⩾10) of pups at weaning.

(**H-I'**) Carmine staining of the wholemount mammary glands (**H**, **I**) and H&E staining of sections (**H'**, **I'**) of the control (**H**, **H'**) and *Pou2f3* knockout (**I**, **I'**) mammary glands during lactation. N =3.

(**J, K**) Whole-mount carmine staining of mammary glands whose cleared fat pads were transplanted with 10,000 basal epithelial cells that were *Pou2f3* heterozygous (**D**) or null (**E**). Scale bars: 5 mm. N = 5.

Data are mean ± SD. Statistical analysis was performed using unpaired Student's t-test. *p < 0.05; **p < 0.01; ***p < 0.001; ****p < 0.0001; n.s., not significant.

**Supplementary Figure 4: POU2F3 Promotes Basal-to-Luminal Conversion by Restricting Chromatin Accessibility to the Basal Transcriptional Program.**

(**A-H**) An in vitro minigland culture assay where control (**B-B'''**) or *Pou2f3*-GOF (**C-D'''**) basal cells were embedded and cultured for 14 days, during which basal organoids underwent luminal differentiation and epithelial branching. (**A**) Measurement of the levels of *Pou2f3*-overexpression based on qPCR reaction. N = 3. (**E**-**H**) Quantification of the effects of *Pou2f3*-GOF on epithelial branching (**E**) and luminal differentiation (**F**-**H**) based on FACS analysis. Note that around half of the *Pou2f3*-GOF growth was unbranched (**E**). N ≥ 5.

(**I**) Heatmap analysis of the top 50 most differentially up- or down-regulated genes in the *Pou2f3* null miniglands when compared with the control miniglands.

(**J, K**) Representative ATAC-seq track plots showing differential accessibility at the sites, underscored by a red line, of *Itga6* (**J**) and *Trp63* (**K**).

**Supplementary Figure 5: *Pou2f3* is Essential for Cell Type Distribution During Mammary Gland Development.**

(**A-D**) Mammary gland development in *Pou2f3* knockout mice. (**A, B**) Whole-mount Carmine staining of mammary glands showing comparable branching morphology between *Pou2f3* wild-type (A) and heterozygous (**B**) mice. (**C, D**) Quantification of ductal elongation (**C**) and branching points (**D**). Scale bars: 2 mm. N ≥ 3 mice.

(**E-J**) Immunofluorescence staining of sections of the mammary epithelium using the indicated luminal (**E-F'**) and epithelial (**G-J'**) cell markers from *Pou2f3* control (**E, G, I**) and knockout (**F, H, J**) mice. Scale bars: 20 µm. N ≥ 3 mice.

(**K**) Quantification of the percentage of luminal cells (K8+, as shown in Fig. 5E-F") in the mammary epithelium.

**(L-N)** FACS analysis of the percentages of basal and luminal cells in the mammary gland of the *Pou2f3* control (**L**) and mutant (**M**) mice. (**N**) Quantification of the percentage of luminal cells in the mammary epithelium. Note a small but significant increase of luminal cells in the *Pou2f3* knockout glands when compared with control glands. N ≥ 3 mice.

(**O**) Quantification of relative cell density in the basal and luminal epithelial layers at the 7-weekstage. N ≥ 3 mice.

**Supplementary Figure 6: A POU2F3-TNF Feedback Loop Maintains Mammary Epithelial Homeostasis.**

(**A**-**F**) Evaluation of the effect of *Pou2f3* loss on the percentages of luminal cells in the mammary epithelium at the 2-week-stage based on immunofluorescence staining of basal and luminal marker expression (**A**-**C**) or flow cytometry (**D**-**F**). N ≥ 3 mice.

(**G**) Quantification of relative cell density in the basal and luminal epithelial layers at the 2-week stage. N ≥ 3 mice.

(**H**) Cell cycle analysis of luminal cells from *Pou2f3* control and mutant mammary epithelium.

(**I, J**) Cell proliferation (**I**) and apoptosis (**J**) assays of *Pou2f3* control and mutant luminal epithelium based on Ki67 immunofluorescence and the TUNEL staining, respectively. N ≥ 3 mice.

(**K**-**M**) Colony-forming assaying using luminal cells derived from the *Pou2f3* control (**K**) and mutant (**L**) mammary glands, and the quantification of colony numbers form by every 500 luminal cells (**M**). N ≥ 6.

Data are mean ± SD. Statistical analysis was performed using unpaired Student's t test. *p < 0.05; **p < 0.01; n.s., not significant. Abbreviations: Rel, relative; expr, expression.

**Supplementary Figure 7: A POU2F3-TNF Feedback Loop Maintains Mammary Epithelial Homeostasis.**

(**A**) GO enrichment analysis of the downregulated genes in *Pou2f3* null basal cells, emphasizing the NFκB signaling pathway.

(**B**) *Tnf* mRNA expression based on scRNA-seq data in luminal cells of the *Pou2f3* control and mutant mammary glands.

(**C**, **D**) *Tnf* mRNA expression in different cell types at the 7-week-stage during development (**C**) and during mammary gland regeneration (**D**) based on scRNA-seq at these stages.

(**E**) KEGG enrichment analysis of the downregulated genes in the intermediate state cells when compared to basal cells during regeneration, emphasizing the TNF signaling pathway and its downstream NFκB signaling pathway.

(**F**) Relative *Pou2f3* mRNA expression, as detected by qPCR, in HC11 cells cultured in medium with the indicated TNF protein concentration. N = 3.

(**G, H**) Relative *Pou2f3* mRNA expression, as detected by qPCR, in HC11 cells cultured in medium with or without TNF protein and inhibitor. (**G**) Effect of the NFκB inhibitor JSH-23. (**H**) Effect of the JNK inhibitor SP600125. N = 3.

**Supplementary Figure 8: Distinct Molecular Programs Govern Cell Fate in Organ Development and Regeneration.**

(**A-D**) Bioinformatics analysis of the cell subtypes, basal and luminal marker expression, and Cytotrace of mammary gland epithelial cells during development.

(**E, F**) Heatmaps of the up- (**E**) or down-regulated (**F**) genes in the P4 luminal intermediate cells during development when compared to embryonic mammary cells and adult luminal cells.

(**G**) GO enrichment analysis of the downregulated genes in the P4 luminal intermediate cells during development when compared to embryonic mammary cells and adult luminal cells.

(**H**) Venn diagrams illustrating the methods by which differentially expressed genes of development and regeneration were extracted.

**Supplementary Figure 9: Distinct Molecular Programs Govern Cell Fate in Organ Development and Regeneration.**

(**A**, **B**) qPCR analysis showing the knockdown efficiency of *Pou2f3* (**A**) and *Egr1* (**B**) using shRNA. N=3.

(**C**, **H**) Effect of *Ehf* and *Egr1* knockdown on basal-to-luminal conversion based on the in vitro minigland regeneration assays. Basal cell aggregates were transfected with lentivirus expressing a scramble control (**C**-**C'**), *Ehf* (**D**-**D'**), or *Egr1* shRNA (**E**-**E'**). N = 3. (**F**-**H**) Flow cytometry analyses of the percentages of basal and luminal cells in the miniglands of control (**F**), *Ehf* (**G**), or *Egr1* shRNA (**H**). N ≥ 6.

(**I**-**Q**) Effect of WNT signaling inhibition using DKK1 protein (**I**-**L**) or the IWR-1 inhibitor (**M**-**Q**), on basal-to-luminal conversion based on the in vitro minigland regeneration assays. Flow cytometry analyses of the percentages of basal and luminal cells in the miniglands of control (**K, O**), DKK1 (**L**), or IWR-1 (**P**). (**Q**) Quantification of the percentage of luminal cells in the organoid epithelium treated with IWR-1. N ≥ 5.

**Supplementary Figure 10: Injury Repair, but not Development, of the Prostate and Pancreas Depends on *Pou2f3* function.**

(**A-E**) Assessment of the percentage of luminal cells based on immunofluorescent in the *Pou2f3* control and null prostate under normal (**A, B''**) or a prostatitis (**C, D''**) condition. (**E**) Quantification of the percentage of luminal cells based on immunofluorescent microscopy. N ≥ 4 mice.

**(F-G')** Immunofluorescence of the luminal marker K19 on a frozen section of *Pou2f3* control (**F**, **F'**) and null (**G**, **G'**) pancreas, showing normal luminal differentiation and epithelial architecture. N ≥ 4 mice.

Data are mean ± SD. Statistical analysis was performed using unpaired Student's t-test. ****p < 0.0001; n.s., not significant.

**Supplementary Figure 11: Model of *Pou2f3* function.**

(**A**) Model diagram of the mutual regulation of POU2F3 and TNF during mammary gland development and regeneration.

**SupFig. 1. Identification of a Novel Intermediate Cell State During Mammary Gland Regeneration.**

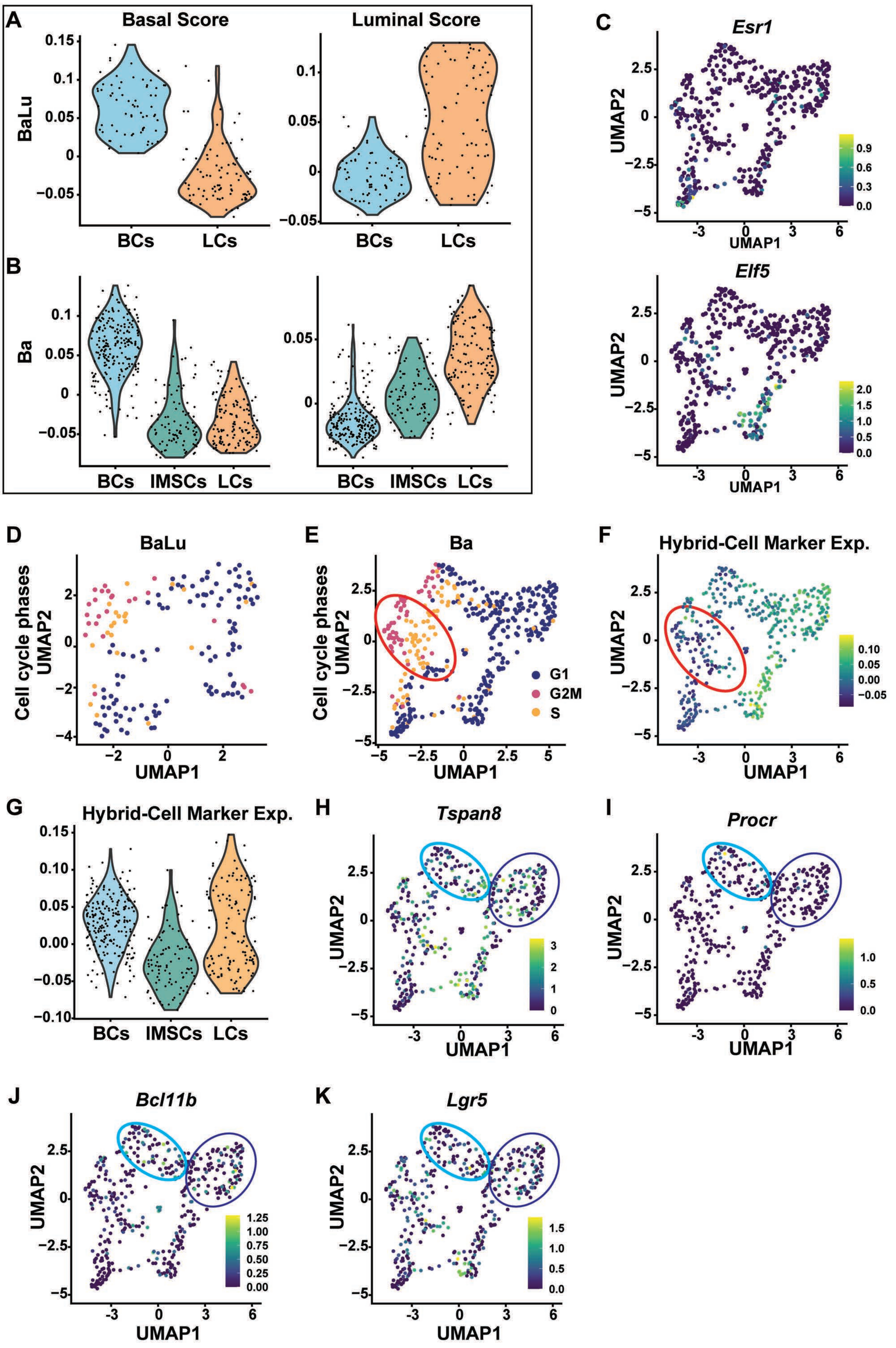

**SupFig. 2. The POU Domain Transcription Factor *Pou2f3* is a Key Regulator of the Intermediate State.**

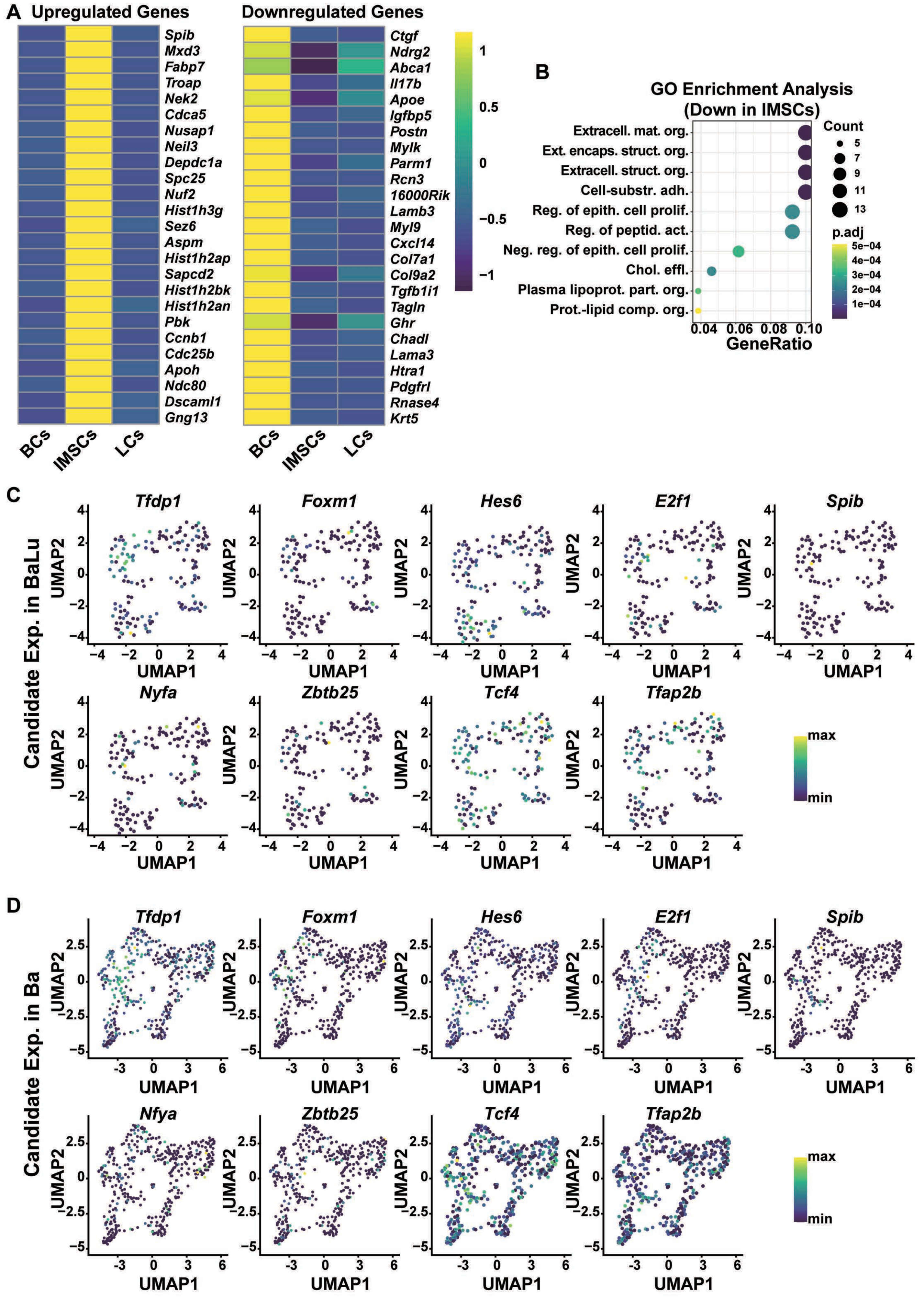

**SupFig. 3. POU2F3 Promotes Basal-to-Luminal Conversion by Restricting Chromatin Accessibility to the Basal Transcriptional Program.**

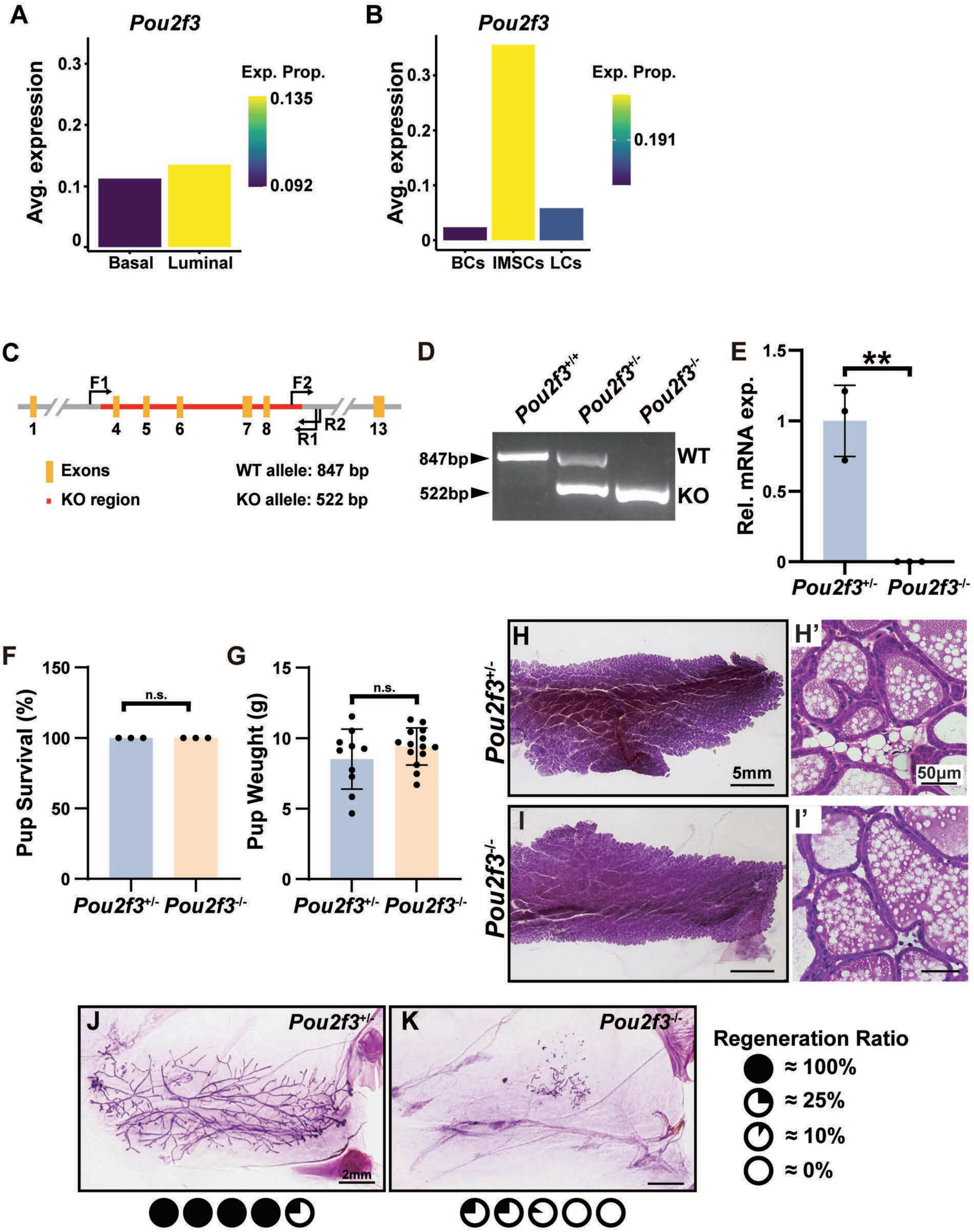

**SupFig. 4. POU2F3 Promotes Basal-to-Luminal Conversion by Restricting Chromatin Accessibility to the Basal Transcriptional Program.**

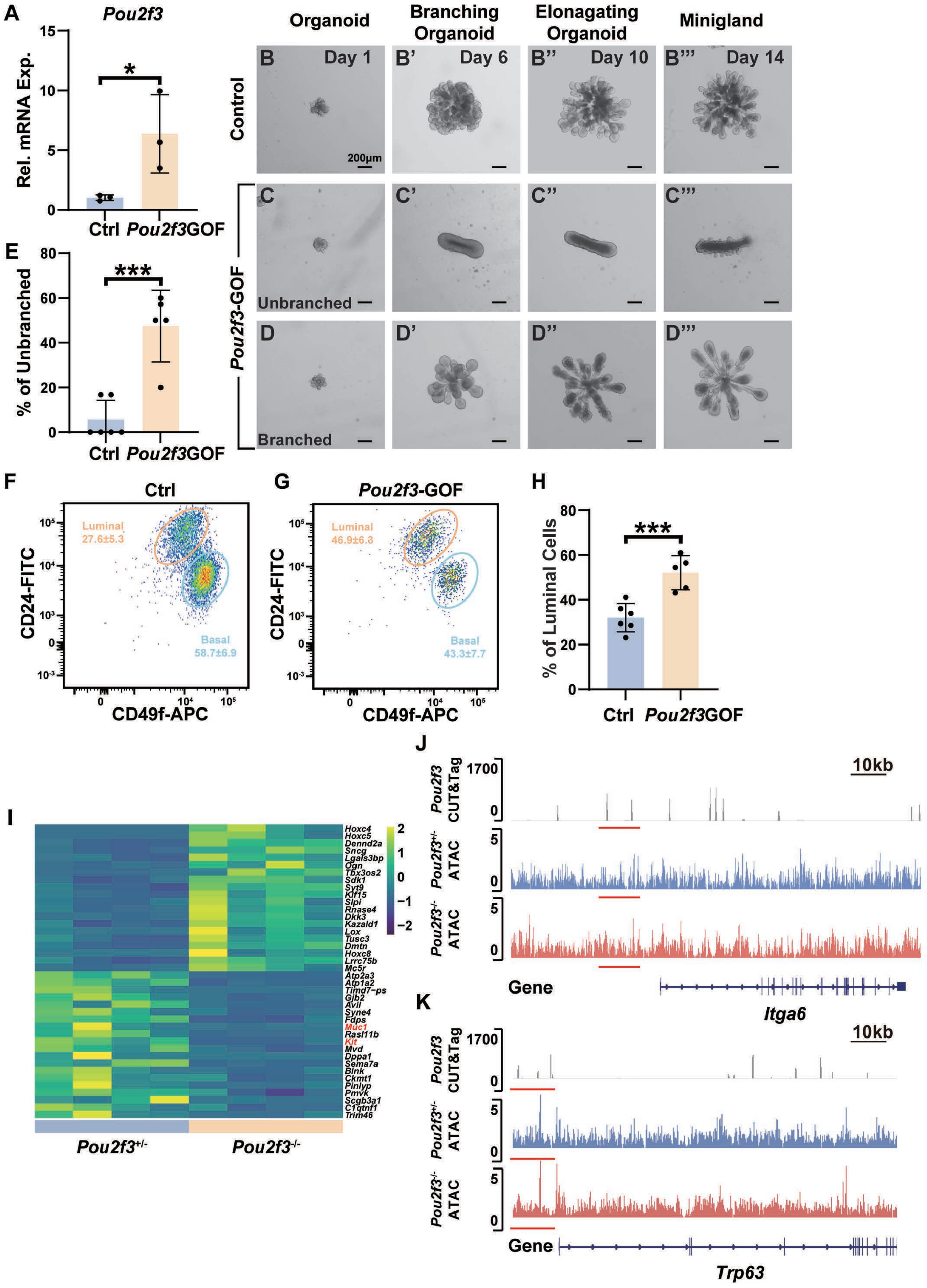

**SupFig. 5. *Pou2f3* is Essential for Cell Type Distribution During Mammary Gland Development.**

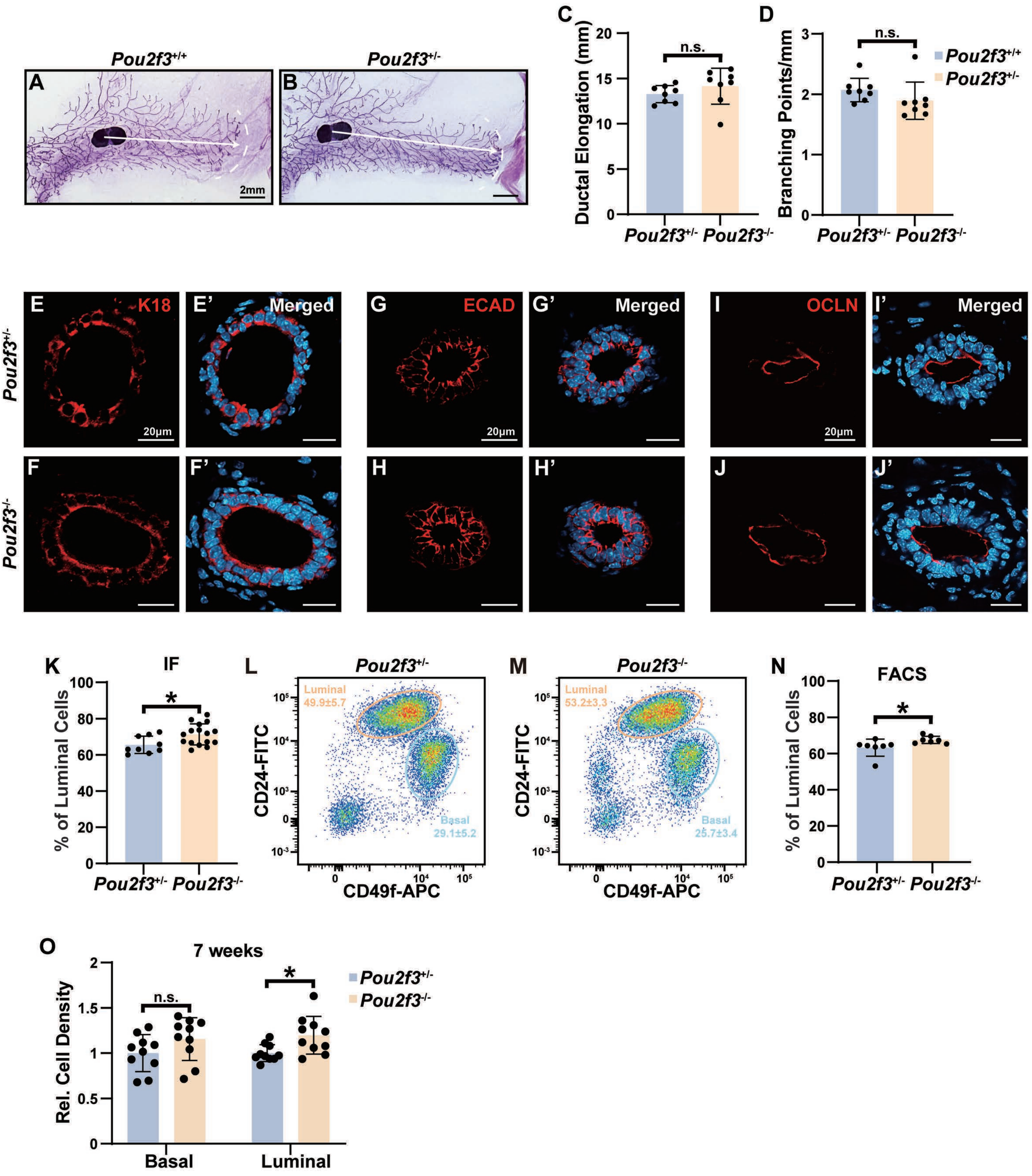

**SupFig. 6. A POU2F3-TNF Feedback Loop Maintains Mammary Epithelial Homeostasis.**

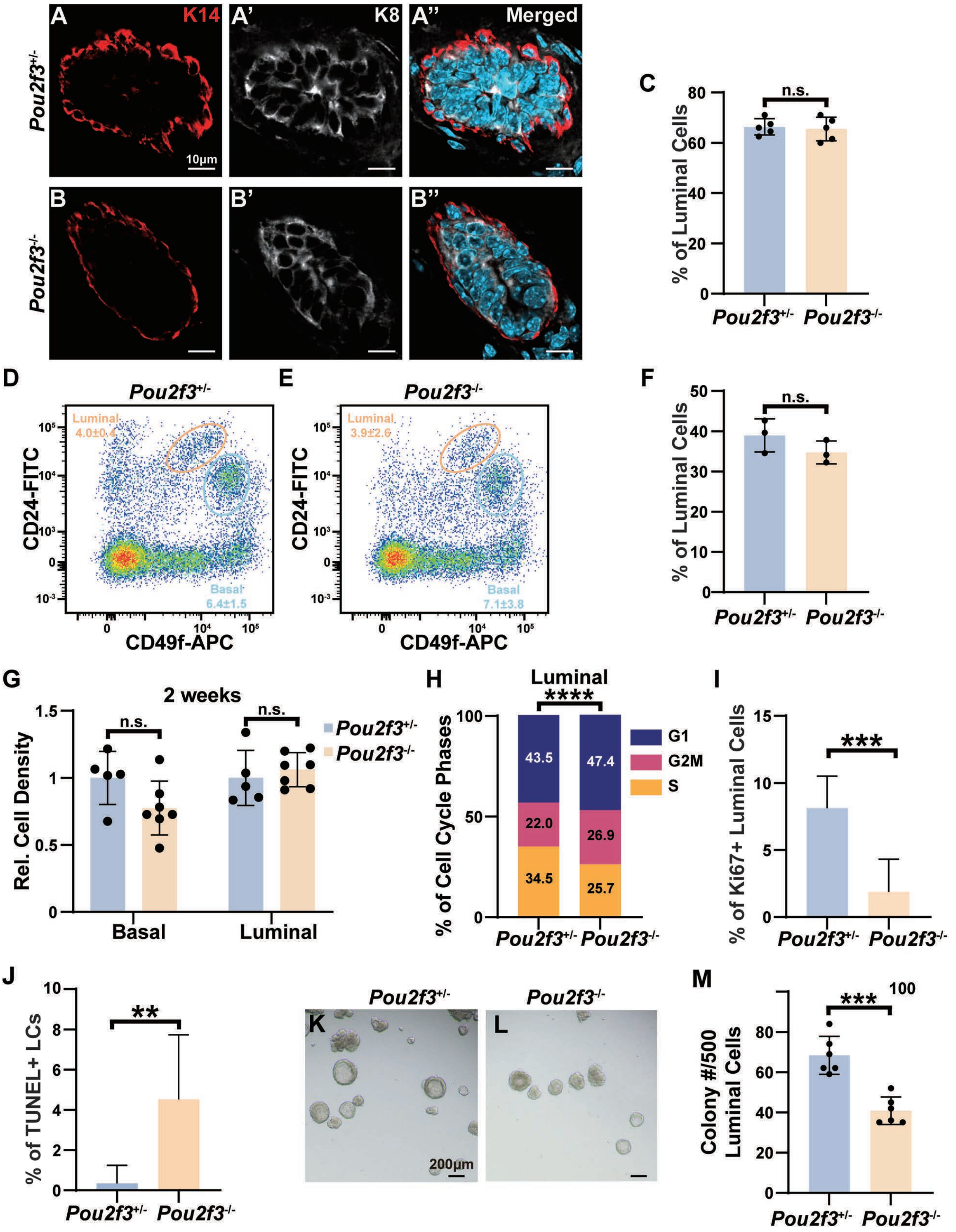

### SupFig. 7. A POU2F3-TNF Feedback Loop Maintains Mammary Epithelial Homeostasis.

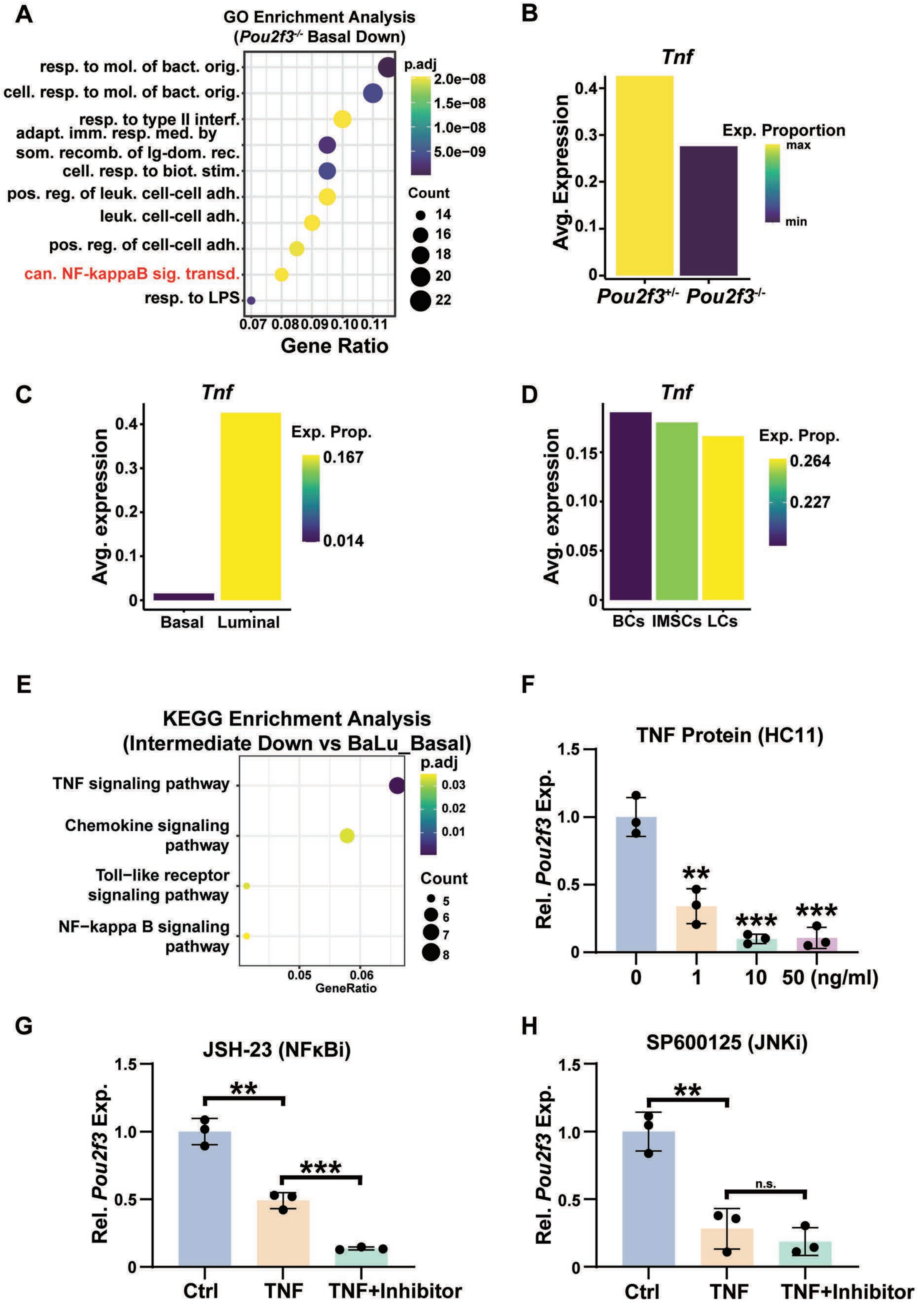

**SupFig. 8. Distinct Molecular Programs Govern Cell Fate in Organ Development and Regeneration.**

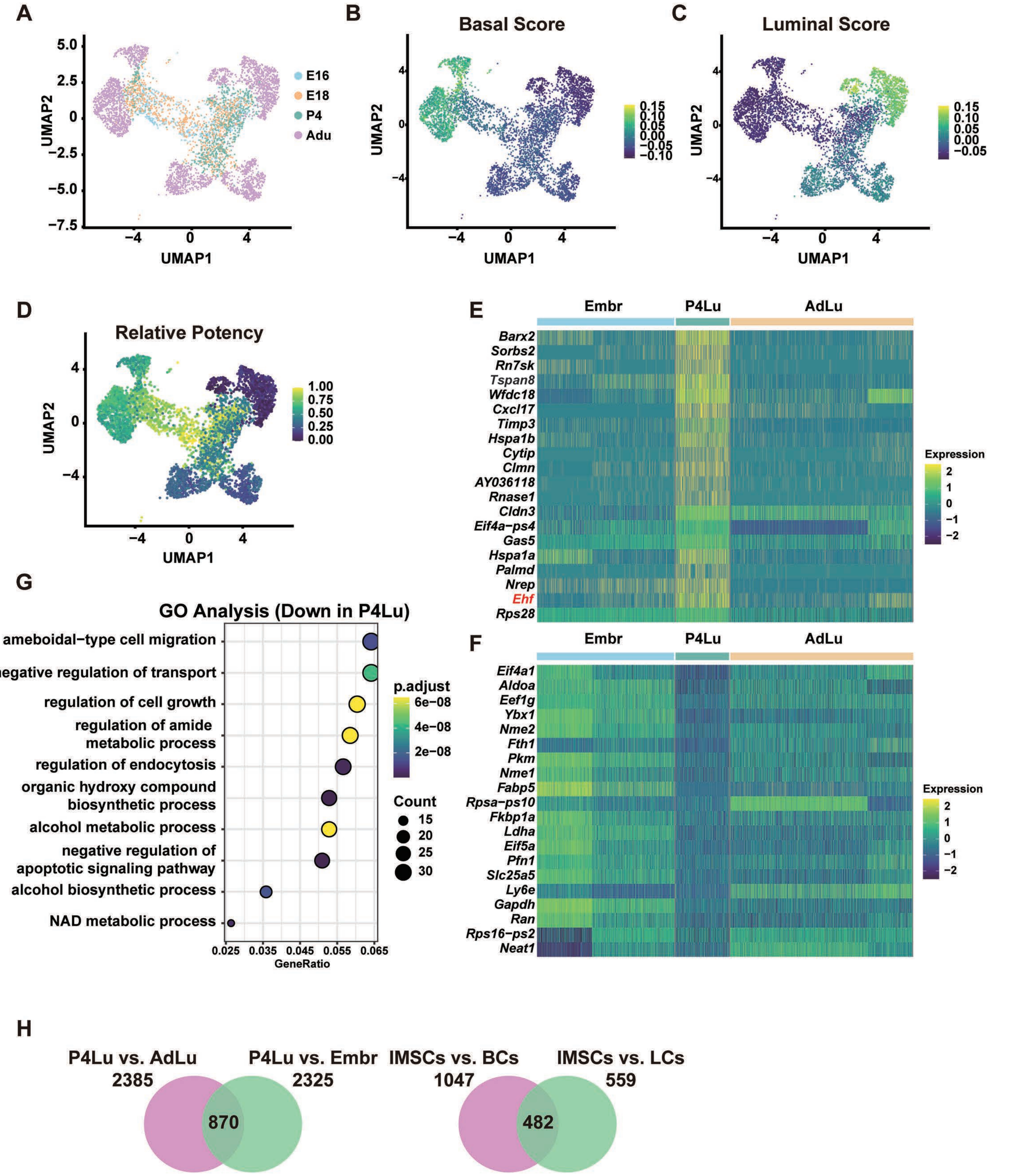

**SupFig. 9. Distinct Molecular Programs Govern Cell Fate in Organ Development and Regeneration.**

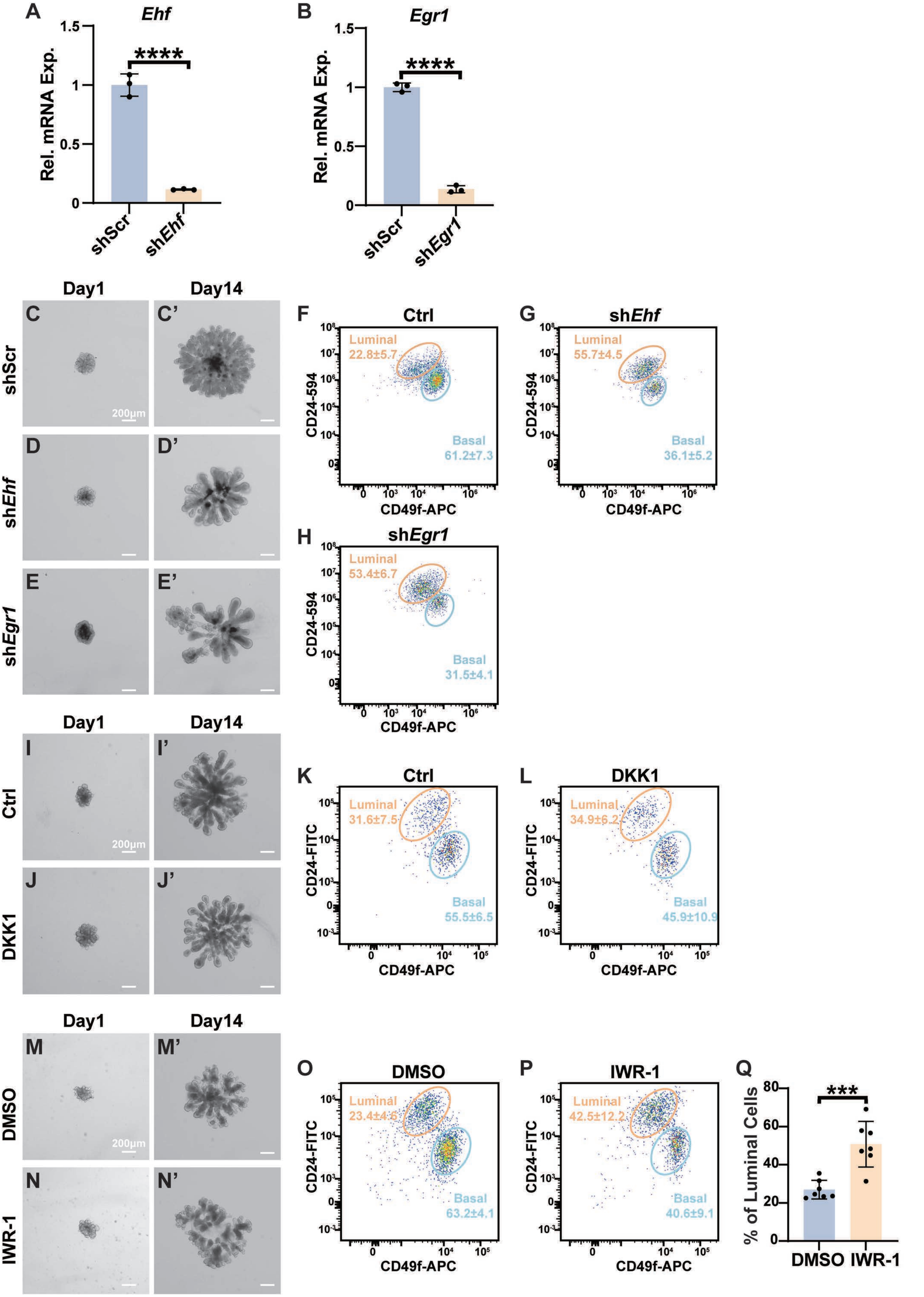

**SupFig. 10. Injury Repair, but not Development, of the Prostate and Pancreas Depends on *Pou2f3* function.**

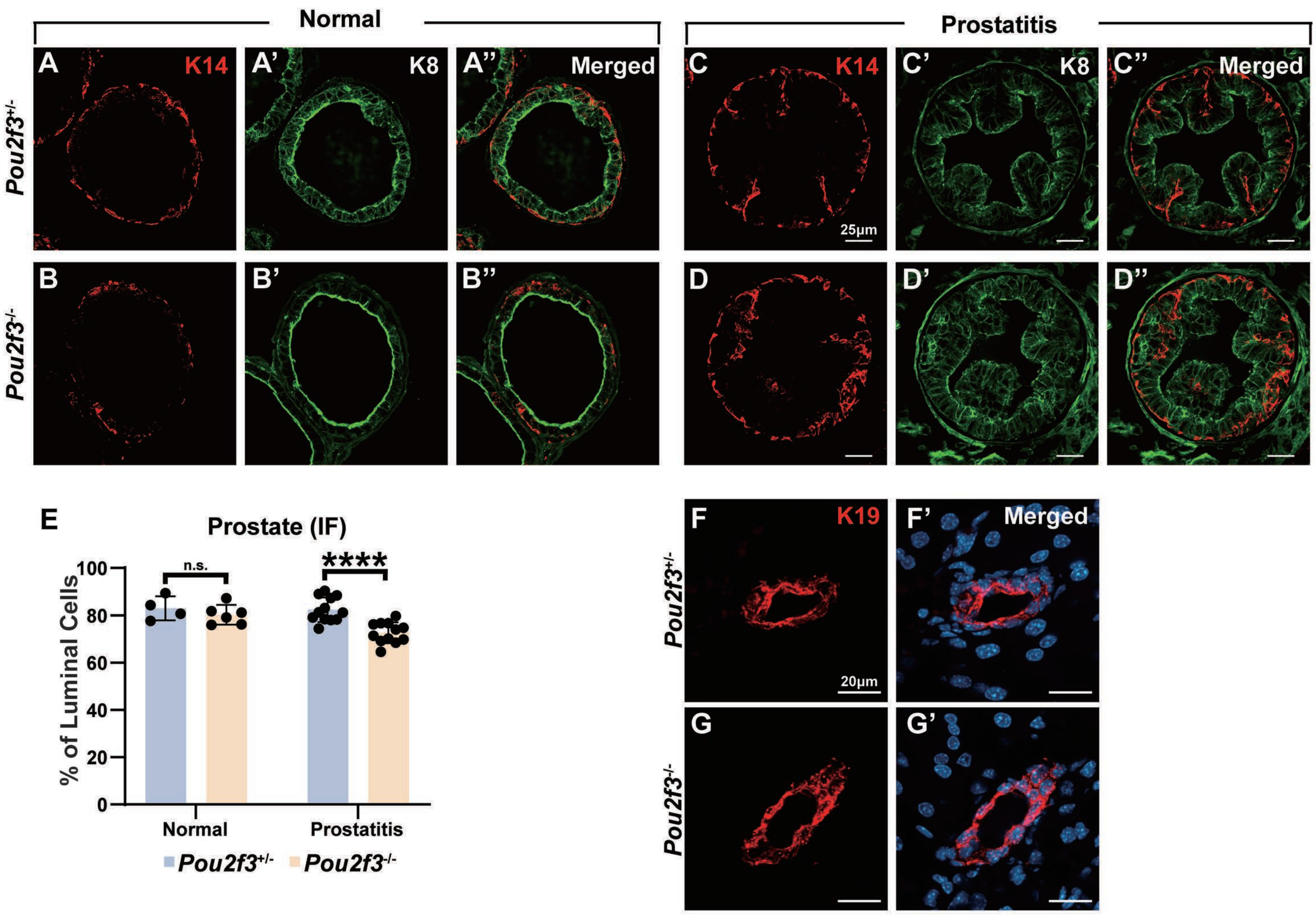

**SupFig. 11. Model of POU2F3 Function.**

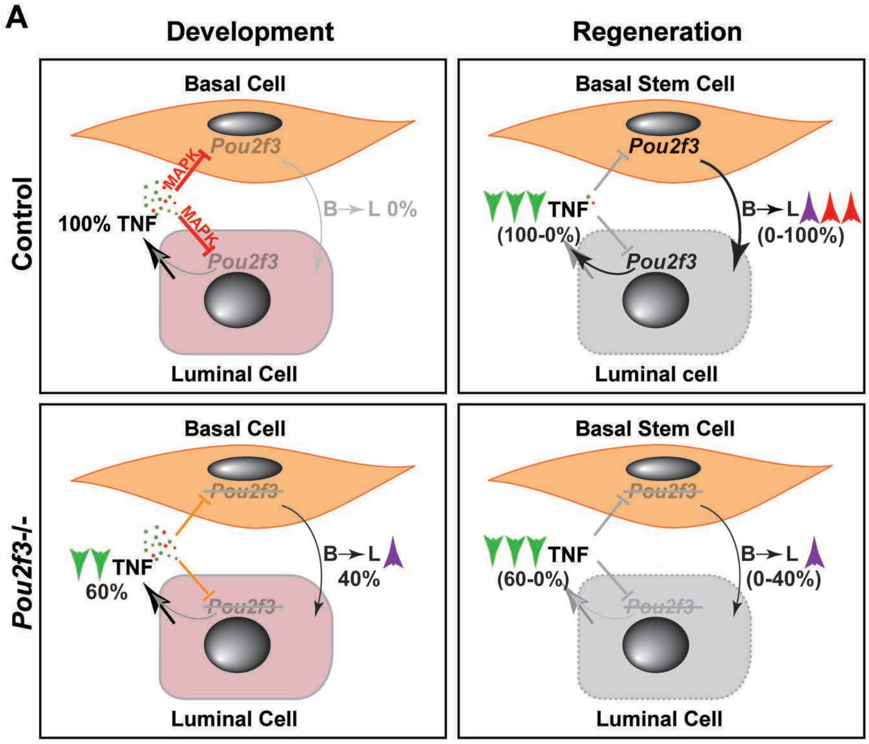
